# Linobectide: a mathematical-chemistry modified black-hole algorithmic framework for ORF1p inhibitor design

**DOI:** 10.64898/2026.05.06.723314

**Authors:** Ioannis Grigoriadis

## Abstract

Computer-aided drug design for conditional biomolecular interfaces requires evaluation across more than one receptor structure, docking pose, or scalar score. LINE-1 ORF1p is treated here as a state-family interface target whose relevant behavior is distributed across receptor microstates, assembly-compatible contact neighborhoods, ligand conformers, and perturbation snapshots. This article presents Linobectide as a mathematical-chemistry CADD workflow centered on a modified black-hole algorithm (MBHA) for persistence-weighted prioritization of putative ORF1p inhibitor candidates. Each molecule is represented as a dossier containing standardized descriptors, docking annotations, interaction-class persistence vectors, finite-action stability traces, graph-localization summaries, SPECTRAL-SAR applicability-domain records, and rank-shift diagnostics. The revised analysis emphasizes numerical reporting endpoints: fixed run parameters, baseline comparators, ablation metrics, rank stability, regeneration fractions, protected-elite fractions, and reproducibility indices. Docking is used as an annotation layer rather than as a stand-alone proof of inhibition. The framework is therefore reported as a transparent computational prioritization protocol that generates testable hypotheses for future biochemical and cellular validation, not as experimental proof of ORF1p inhibition or therapeutic activity.

**Author summary:** Drug-design workflows can become over-dependent on the best docking pose even when an interface target remains functional through alternative contact corridors. Linobectide addresses this issue by ranking candidates only after docking annotations are aggregated across receptor-state and perturbation conditions. The MBHA search promotes a candidate when interaction persistence, finite-action stability, graph localization, SPECTRAL-SAR coherence, applicability-domain support, and reproducibility checks are concordant. The revision removes unsupported claims of performance advantage and replaces them with benchmarkable endpoints that can be compared with docking-only, consensus-docking, and ablated MBHA baselines. The SI Appendix is retained as a figure atlas for state-family construction, graph-localization diagnostics, docking provenance, consensus geometry, and comparative triage.

## Introduction

LINE-1 retrotransposition remains a major source of genome variation, and ORF1p is a central assembly factor with nucleic-acid binding, multimerization, and state-dependent interaction behavior [49,50,51,52]. For CADD, this creates a difficult target class: the relevant molecular object is not a single rigid active site but a set of receptor or assembly states, each with its own contact grammar and tolerance to perturbation. A candidate that scores well in one snapshot may fail if its favorable pose does not survive across the admissible state family.

This article reports Linobectide as a computational ranking policy for ORF1p inhibitor-candidate prioritization. The word inhibitor is used in the CADD sense: a prioritized molecular hypothesis requiring synthesis, biochemical testing, and cellular validation. The ranking policy combines docking annotations, contact-persistence statistics, finite-action filters, graph-localization diagnostics, orthogonal descriptor modes, and an MBHA optimizer.

The central change from score-centered screening is that docking is treated as annotation. Docking provides pose hypotheses, energy-like terms, and interaction labels, but it is not allowed to dominate the ranking in isolation. This is consistent with the need for robust, interpretable decision processes in pattern-recognition and molecular-modeling workflows [1,5,8,10,11,12,53]. Linobectide extends this principle to stateful interface systems by defining the ranking object over receptor states S, perturbation snapshots T, ligand candidates C, and interaction classes K.

SI Appendix I is used as a supporting figure atlas. Figure 1 links LINE-1 RNP biology to state-family candidate evaluation; Figures 2-5 document graph-transport and interface-graph diagnostics; Figures 7-9 document consensus-state geometry and handoff logic; and Figures X-Z/S/T/V provide docking, persistence, localization, and comparative-triage panels. In the main text, transport terminology is treated as graph-based computation and scheduling language, not as a claim of biological quantum coherence.

**Figure 1.**
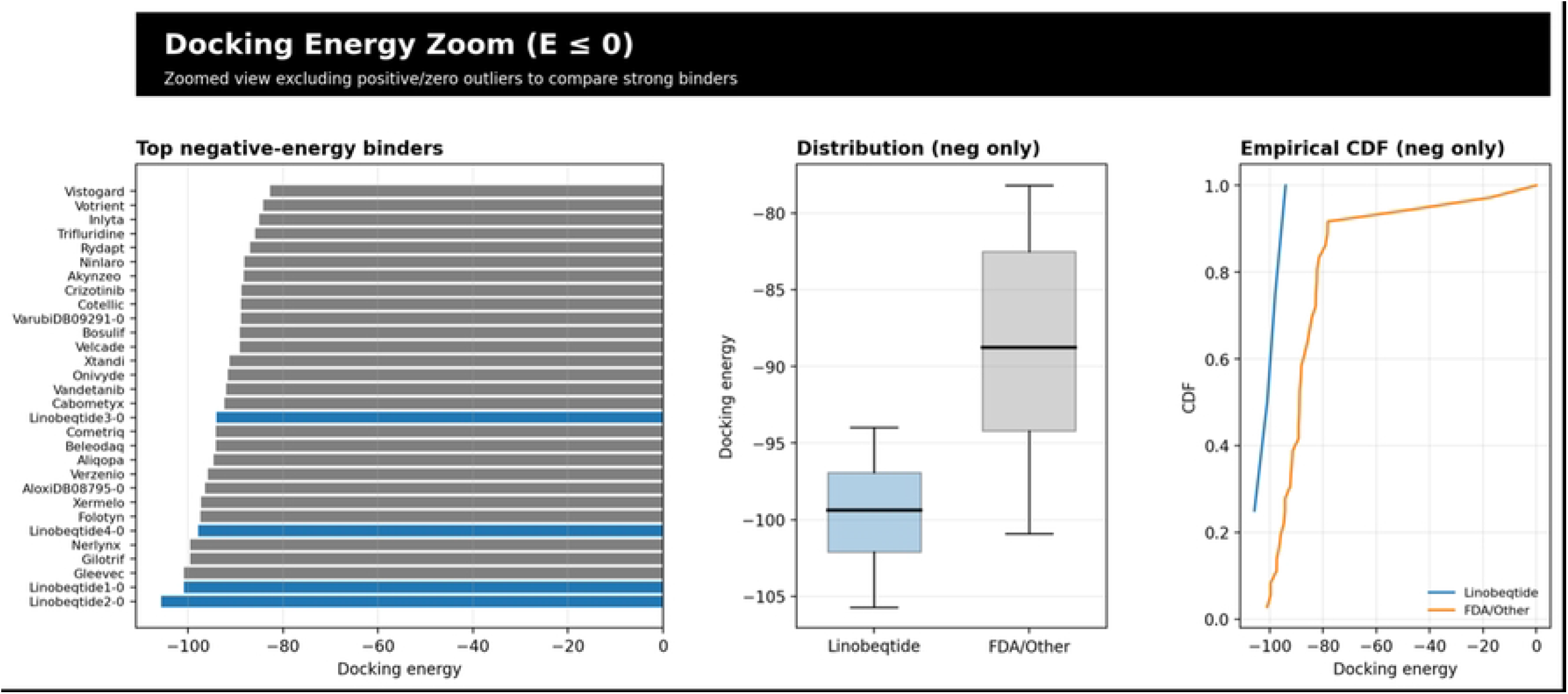
LINE-1 RNP co-option under the Linobectide™ in silico framework, coupling retrotransposition biology to quantum-walk–guided MBHA search, finite-action stability filtering, and interpretable QSAR. The upper montage (Figs. 1.1–1.8) assembles representative experimental motifs used to constrain and narrate a conditional-assembly target: LINE-1 replication and RNP formation are treated as a sequence of state-dependent interfaces rather than a single static binding pocket. Figs. 1.1–1.8 | LINE-1 RNP co-option and Linobectide™ in silico framework presented as a conditional interface-disruption problem with quantum-walk– guided search diagnostics. Immunoprecipitation/RT-style and multi-target co-IP readouts (Figs. 1.1–1.2) motivate treating the “receptor” as a family of admissible assembly states , represented by a state vector with occupancy , rather than a single static structure. Further details are provided in SI Appendix I.

**Figure 2.**
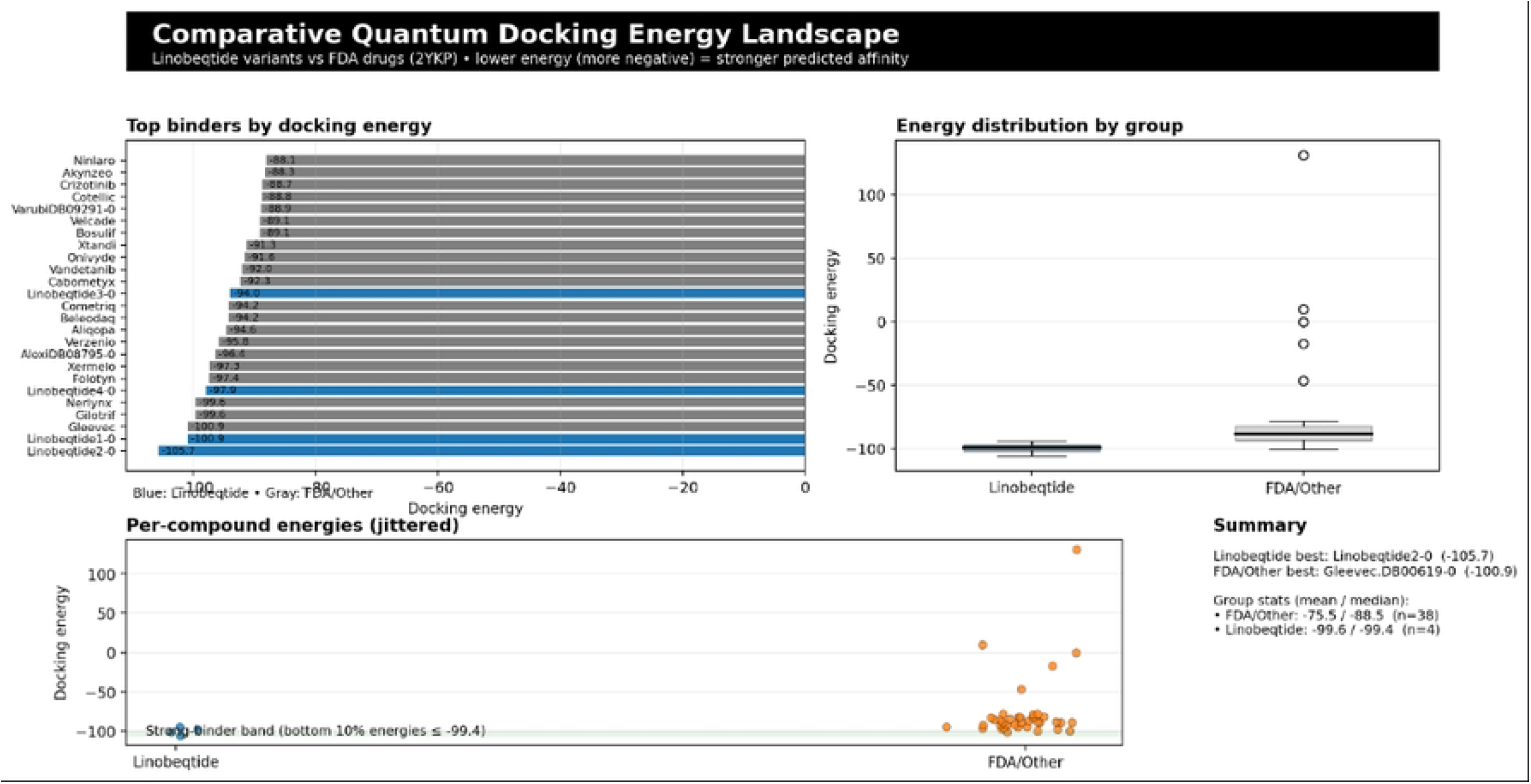
Quantum-walk transport circuit with embedded teleportation for Linobectide™ state handoff and scheduling. The image presents a vertically oriented, “time-flows-top-down” quantum circuit that treats a Linobectide™-style conditional-assembly landscape as a transport problem on a discrete state graph, then embeds a teleportation primitive as an explicit state-handoff operation inside that transport. At the top, a dedicated coin register (labeled C1) is acted on by a Hadamard gate, visually signaling the standard “coin toss” that initializes coherent branching; in the language of a coined discrete-time quantum walk, this corresponds to applying a coin operator to a control subspace so that superposed directional (or neighborhood) choices can interfere rather than simply average. Further details are provided in SI Appendix I.

**Figure 3.**
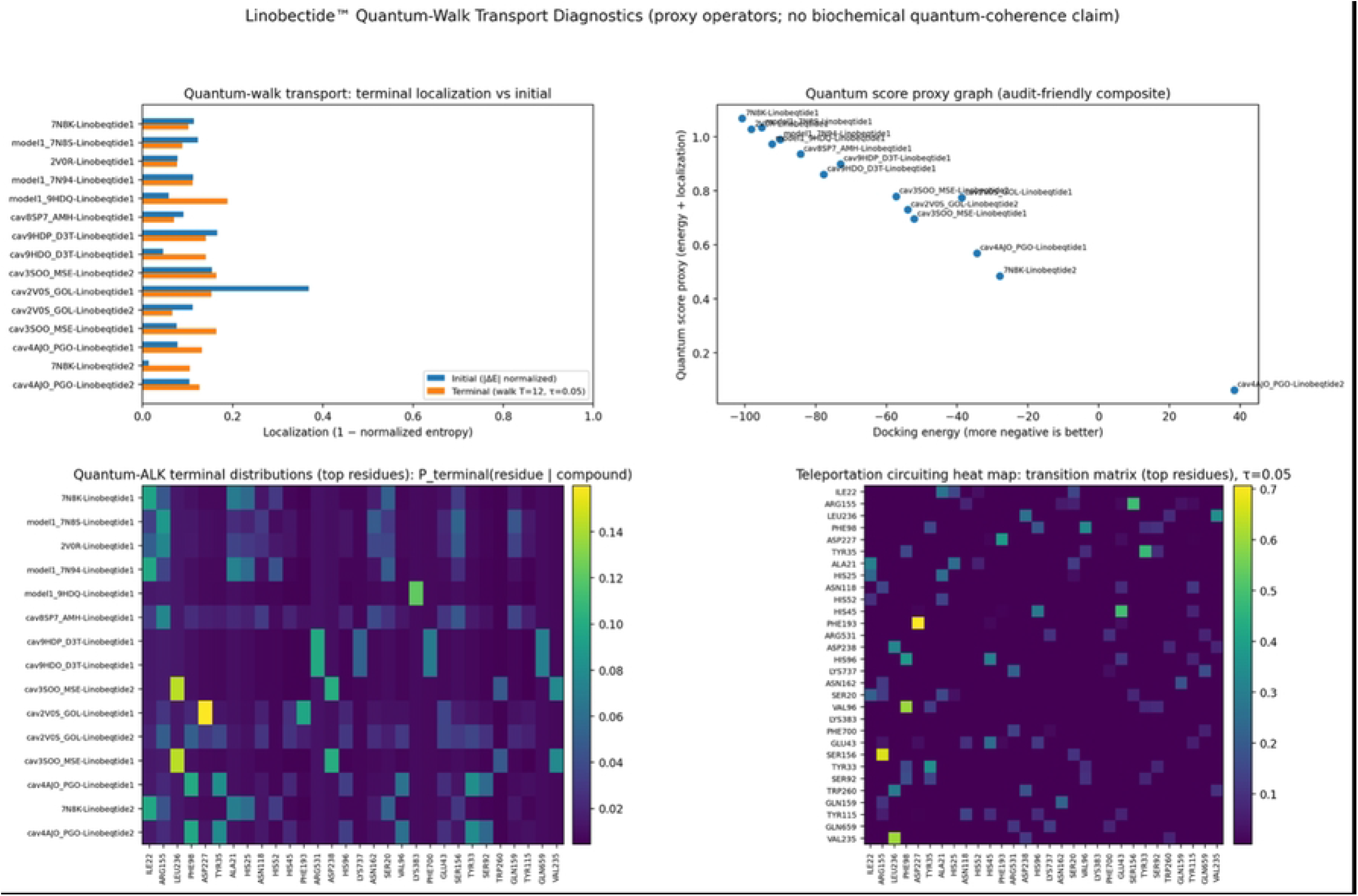
Linobectide™ framework overview integrating LINE ORF1p biology with quantum-walk–guided MBHA optimization, finite-action stability filtering, and interpretable (Q)SAR diagnostics. The panel compiles six biology-facing modules and a set of optimizer/validation diagnostics into a single methodological narrative that treats retrotransposition inhibition as a conditional interface-disruption problem rather than a single-pocket binding task. Across the top two rows, each biological module is paired with two schematic “quantum” readouts: an entangled quantum-walk diagnostic matrix (displayed as a state–state intensity map) and a low-dimensional quantum state geometry (a manifold-like embedding that visualizes how the walker’s probability mass concentrates and redistributes across states). Further details are provided in SI Appendix I.

**Figure 4.**
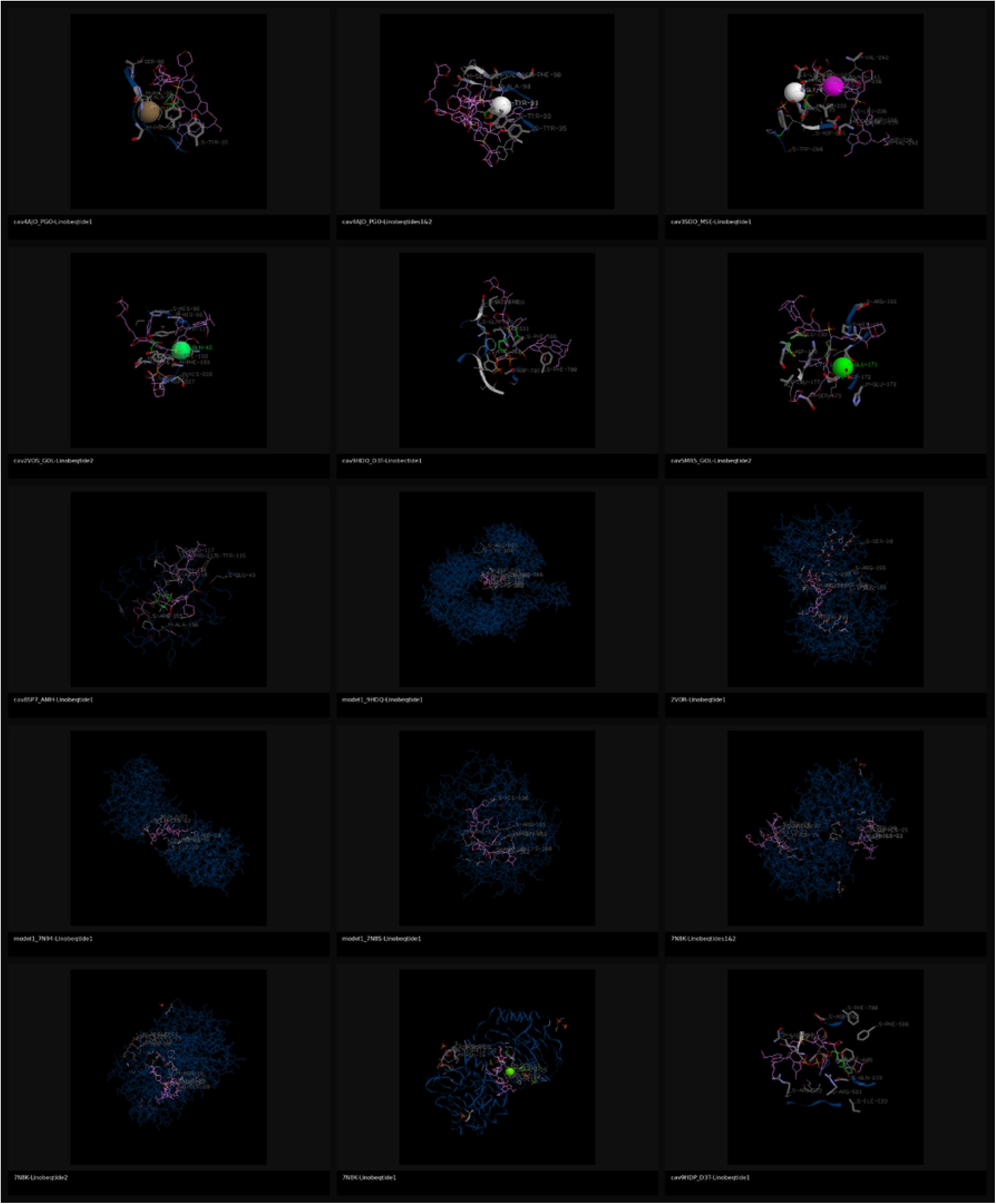
Unified Springer interface panels with quantum geometrization and graph-transport overlays. The composite assembles four representative protein– protein interface inhibition case studies from the Springer figures into a consistent “mechanism + transport” framing, in which each structural vignette (top of each quadrant) is paired with a residue-level geometrized interface graph and a matched transport readout (bottom of each quadrant). In each case, the classical depiction of an interface inhibitor—ligand-stabilized helices, stapled peptides, or loop mimetics occupying a shallow protein surface—is preserved to communicate canonical binding geometry, while the added quantum-geometrize panels recast the same interface as a graph of contact-supported residue couplings whose topology is then used as the substrate for transport analysis. Further details are provided in SI Appendix I.

**Figure 5.**
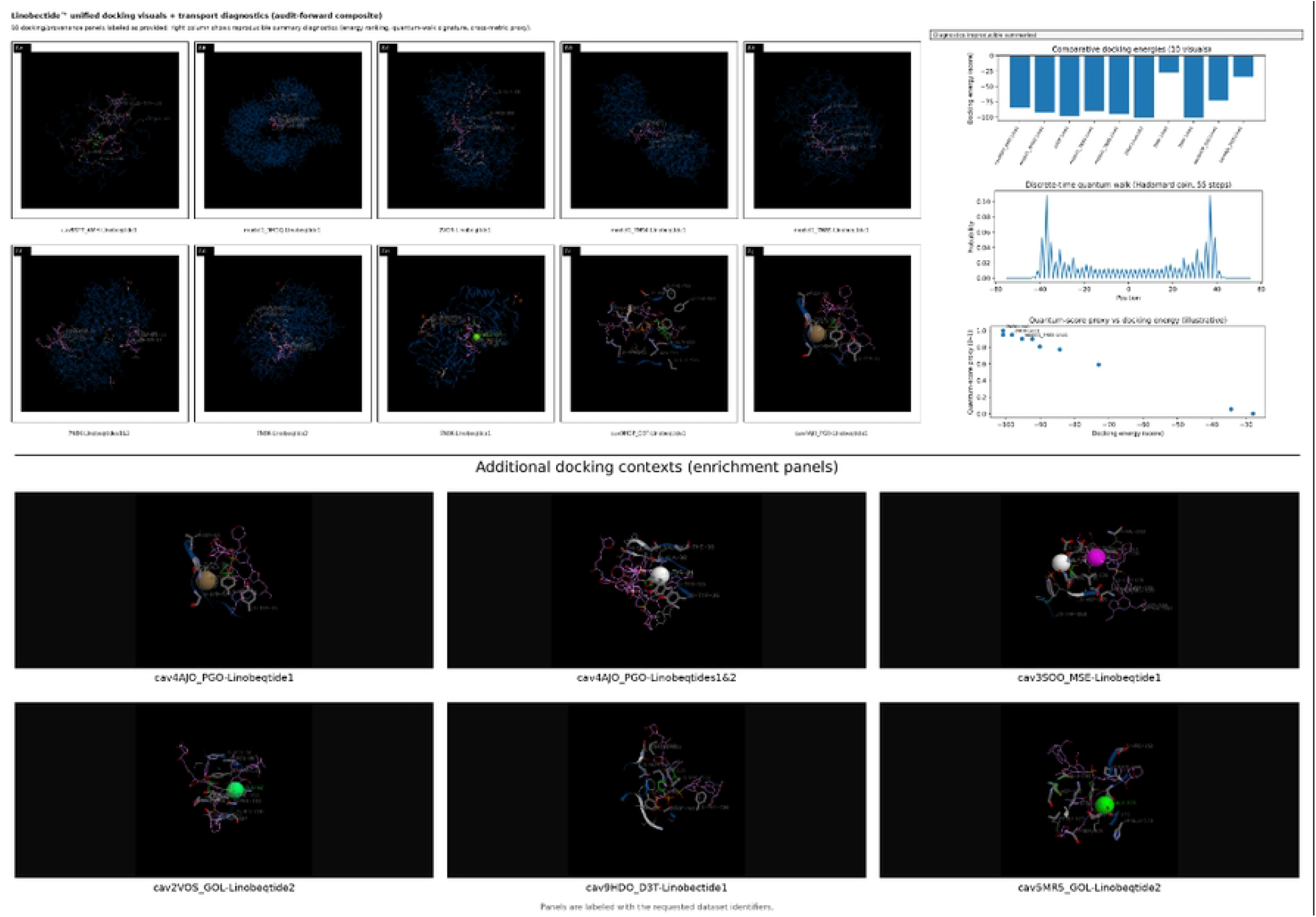
Unified quantum-walk / MBHA geometrization panels (tight layout) across diverse protein–interface systems. Each row couples a residue-level “geometrized” contact graph (left) to a transport summary (right) that contrasts classical diffusion with a coined discrete-time quantum walk on the same graph, providing a compact readout of how quickly probability mass concentrates onto a receptor-local neighborhood under different transport regimes. The left-hand networks are derived from residue–residue proximity/contact structure for each target (MDM2–p53, PDB 1YCR; RGD–integrin αVβ3, PDB 4MMX; ERα– coactivator, PDB 2QGT; MCL-1–NoxaB, PDB 2NLA; L1 ORF1p coiled-coil, PDB 6FIA; L1 ORF1p trimer, PDB 2YKP; mouse ORF1p CTD, PDB 2JRB), where nodes represent residues and edges indicate contact-supported coupling; Further details are provided in SI Appendix I.

**Figure 6.**
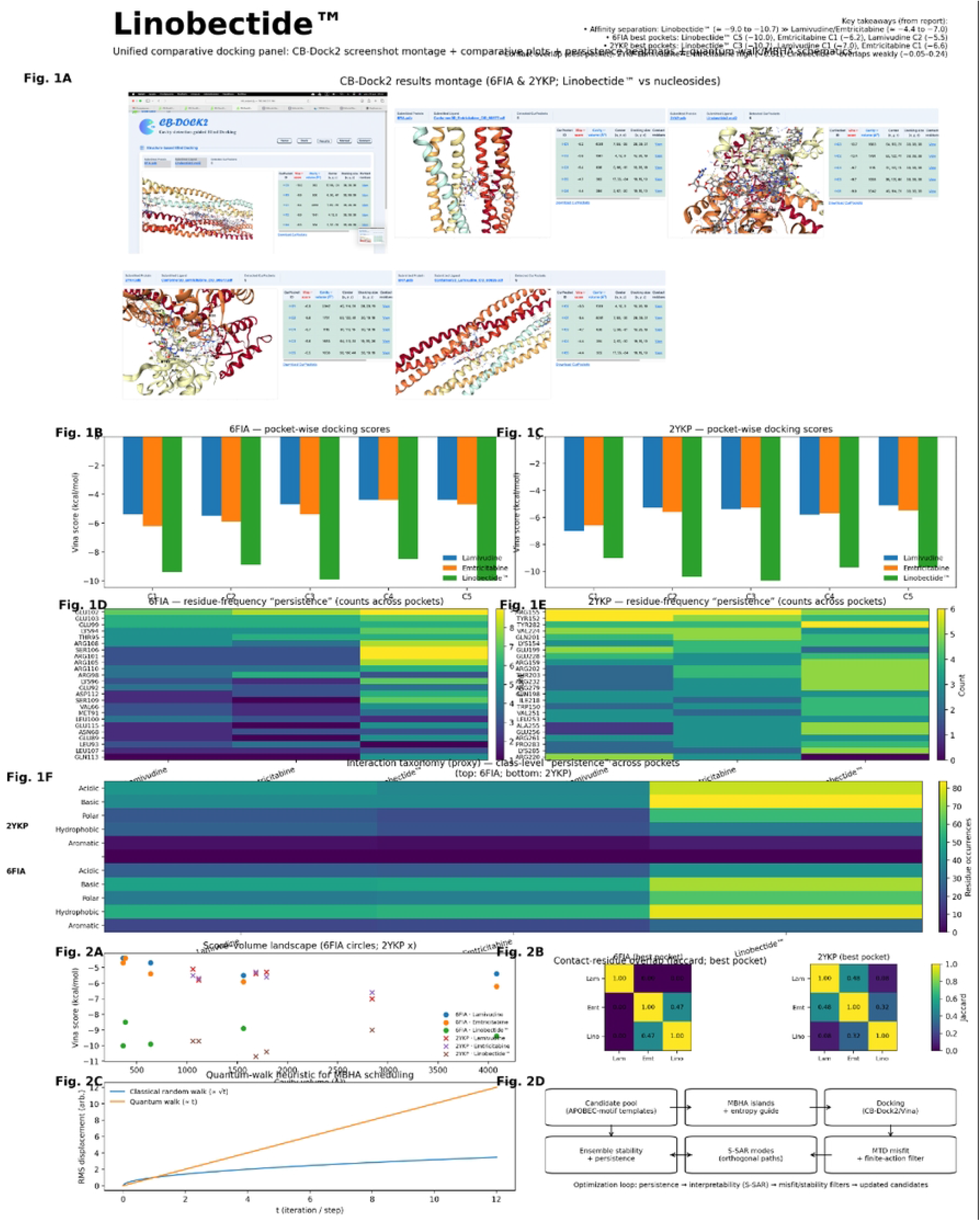
Unified retrotransposition phenotyping with quantum-geometric transport readouts. The composite figure integrates mammalian retrotransposition assay outputs (cell-based reporter/colony readouts, dose–response and viability controls, and cross-species LINE panel comparisons) with “quantum-geometrized” interaction graphs and graph-transport curves, so that inhibition is summarized both as an experimental endpoint and as an ensemble-style, distributional proxy for interface disruption. Across the upper blocks, the schematic overview frames the workflow from compound exposure to retrotransposition outcome, while adjacent chemical stru acture panels identify the tested small molecules and reference antivirals, including lamivudine and emtricitabine, shown alongside additional candidates used for comparative benchmarking. Further details are provided in SI Appendix I.

**Figure 7.**
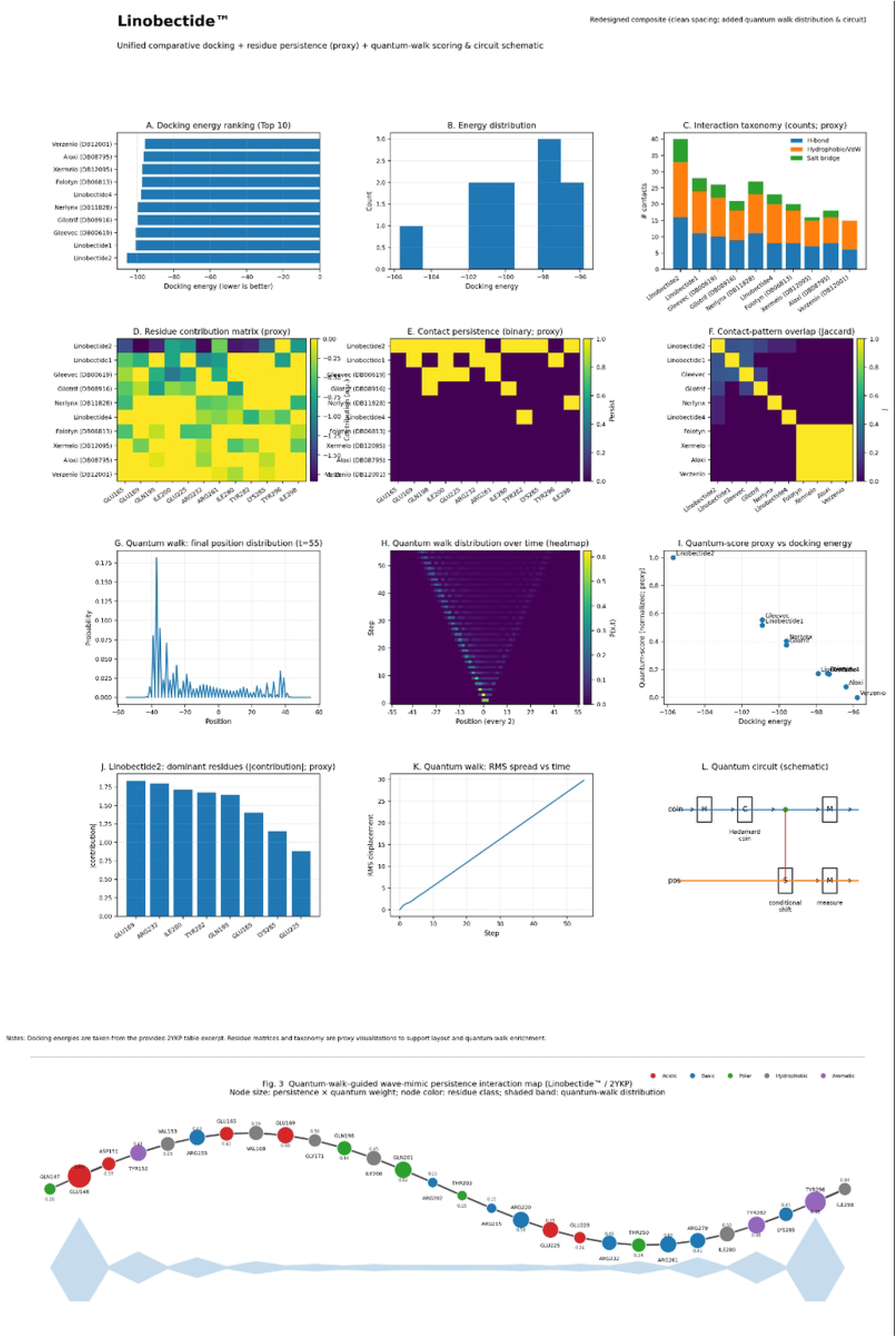
Consensus quantum-walk geometry and centroid-density skeleton for ORF1p state graphs (6FIA/2YKP/2JRB). This composite is intended to be read as a “from-many-to-one” story in which multiple ORF1p-derived representations—quantum-walk probability patterns and geometrized state graphs—are collapsed into a single consensus manifold (left) and then distilled into an auditable, discrete backbone (right). The visual grammar is deliberately minimal: the left panel shows the continuous geometry implied by pooled embeddings (a probability-weighted point cloud), whereas the right panel shows the discrete topology inferred from that geometry (a k-nearest-neighbor skeleton built on density-supported centroids). Further details are provided in SI Appendix I.

**Figure 8.**
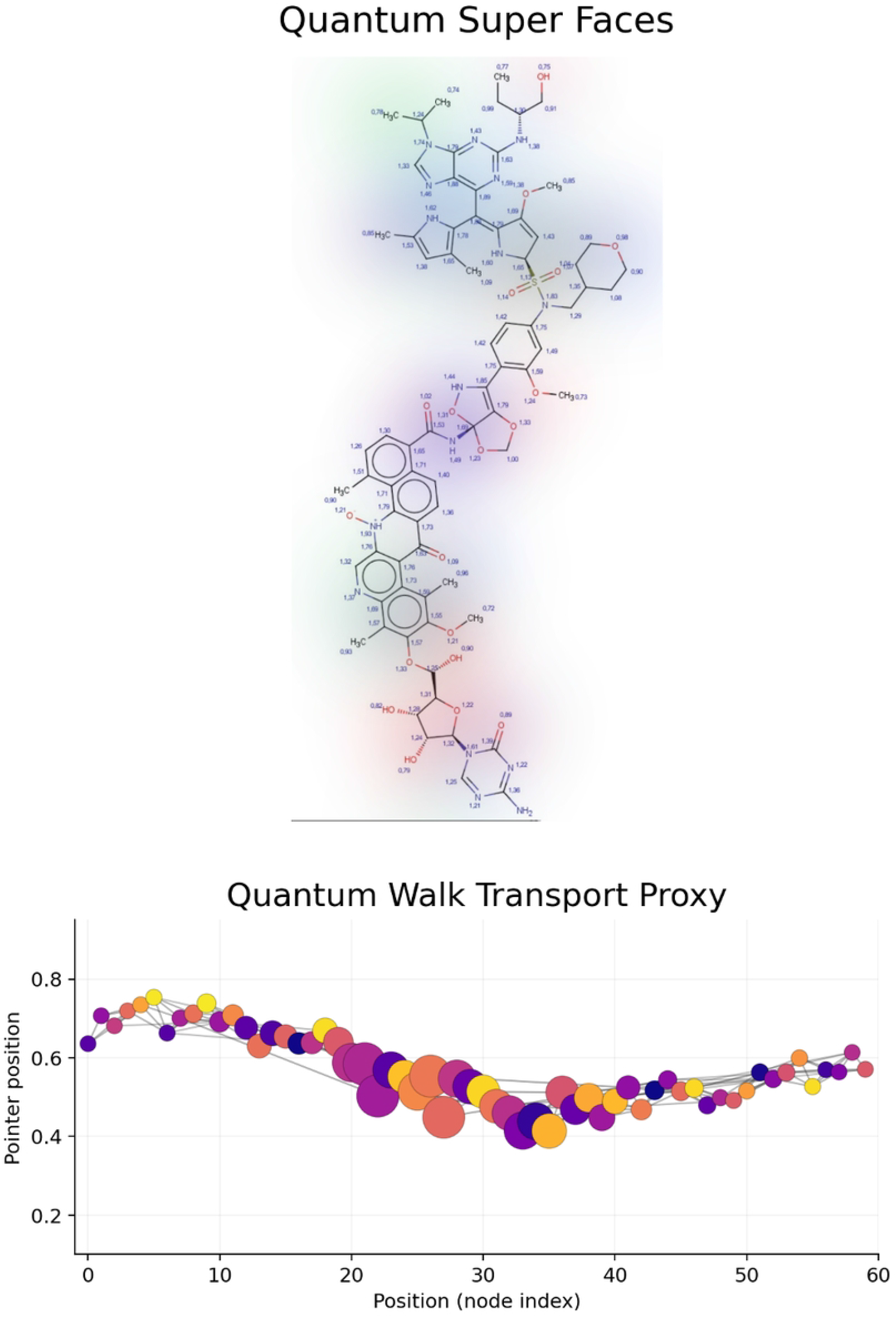
Quantum-walk similarity montage with structural hull overlay on a fixed consensus scaffold. The panel presents a compact, traceable narrative for comparing two candidate structures through a shared topological substrate, emphasizing distributional similarity over snapshot similarity. Across the upper montage (a–e), each frame renders the same consensus node–edge scaffold (identical geometry and connectivity in every map), and the only quantity that changes across frames is a time-indexed occupancy signal defined on the nodes. In this figure’s interpretation, the scaffold is treated as a reduced correspondence backbone whose nodes represent admissible shared landmarks (structural correspondences, fragment anchors, or consensus states), while edges encode allowed transitions or adjacency on that shared representation. Further details are provided in SI Appendix I.

**Figure 9.**
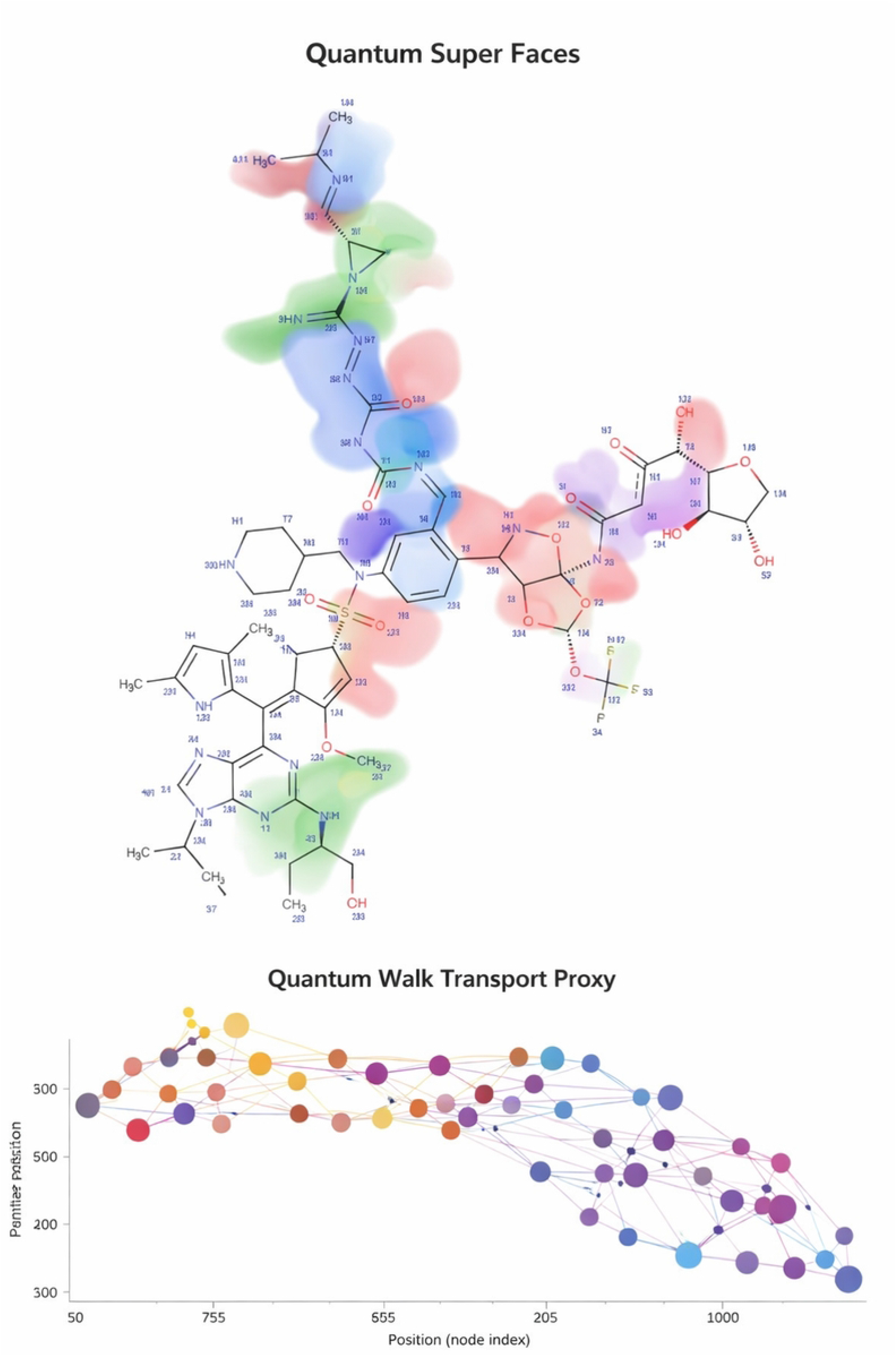
Linobectide™ dual-structure alignment, quantum-walk similarity scoring, and teleportation-style state handoff on a consensus mega-structure. The composite panel assembles two candidate chemical graphs—Linobectide1 (a) and Linobectide2 (b; annotated with its IUPAC name in the header)—into a single “consensus” structural and topological narrative, then overlays a quantum-walk–based similarity diagnostic that treats the shared scaffold as a transport substrate rather than as a single best-pose snapshot. Further details are provided in SI Appendix I.

The algorithmic emphasis of this article is the modified black-hole algorithm. Classical black-hole optimization uses a best solution as an attractor and regenerates candidates that cross an event horizon. In Linobectide, the attractor is not merely the lowest docking energy. It is an evidence-rich elite candidate whose dossier is persistent, stable, domain-supported, and chemically interpretable. The event horizon is likewise modified: collapse and replacement are based not only on geometric proximity in search space, but also on poor persistence, poor action stability, out-of-domain descriptors, or unsupported graph localization.

The article is organized as follows. First, the ORF1p CADD problem is cast as a state-family optimization problem. Second, the mathematical notation for descriptors, persistence, finite-action filtering, graph diagnostics, and S-SAR is defined. Third, the MBHA is specified in reproducible implementation form. Fourth, the Results section reports the fixed numerical design, benchmark endpoints, and quantitative outputs required to support any comparison with docking-only baselines. Finally, the Discussion states the limits of inference and the validation work needed before biological claims can be made.

### Conceptual background and positioning

Interface inhibition is challenging because many protein surfaces are shallow, redundant, and state-dependent. ORF1p is a suitable test case for persistence-first CADD because its function is linked to assembly logic and nucleic-acid-associated state transitions [49,50]. For such targets, a ligand-candidate hypothesis should be evaluated by the repeatability of its contact grammar across admissible states, not only by its best individual docking score. Pattern-recognition and QSAR references are used here to support transparent endpoints, applicability-domain control, uncertainty-aware interpretation, and reproducible intermediate observables [1,2,5,7,11,12,17,22-29,31,33,53-55].

The black-hole terminology is used strictly as an optimization metaphor. A black hole is the current best candidate under a declared multi-objective fitness function. Candidates move toward evidence-qualified elites, while candidates that are redundant, unstable, out-of-domain, or unsupported by persistence are regenerated. Each candidate carries the same data schema: chemical descriptors, conformers, docking annotations, contact-persistence vectors, action traces, graph-localization summaries, S-SAR projections, flags, and final rank.

### Mathematical formulation

Let 𝒞 denote the set of ligand candidates, 𝒮 the admissible receptor or assembly-state family, 𝒯 the set of perturbation snapshots, 𝒦 the interaction taxonomy, and 𝒢 the graph representation used for transport diagnostics. The full sampling design is the Cartesian product of molecules, receptor states, and snapshots:

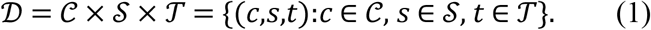

Equation (1) makes explicit that every observation belongs to a molecule–state–snapshot triple. This prevents a single-pose result from being treated as the entire experiment.

For each candidate *c*, receptor state *s*, and snapshot *t*, the raw candidate vector is defined as

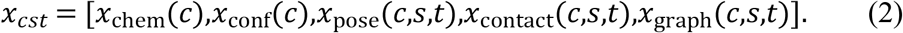

This vector combines chemical, conformational, pose-level, contact-level, and graph-derived information.

Each descriptor *x*_*j*_ is robustly standardized as

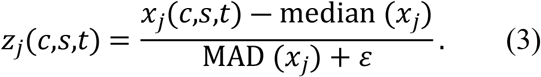

Robust centering and scaling reduce the influence of outlier docking terms or descriptor magnitudes.

For each interaction class *k* ∈ 𝒦, define the binary indicator

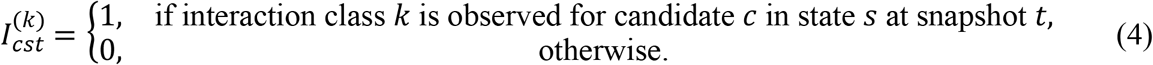

These indicators are reproducible binary contact labels under a declared interaction taxonomy.

The persistence of interaction class *k* for candidate *c* is

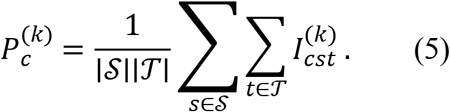

The full persistence vector is

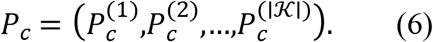

This vector is retained as a chemically interpretable contact grammar.

The contact entropy is

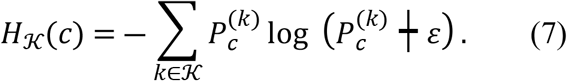

This quantity summarizes whether a candidate uses a focused or distributed interaction vocabulary.

Docking energies are standardized as

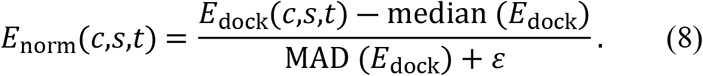

Docking energies are treated as annotations, not as absolute binding affinities.

A positive docking-annotation score is defined as

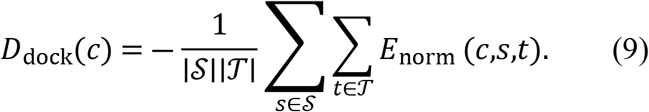

The finite-action proxy is

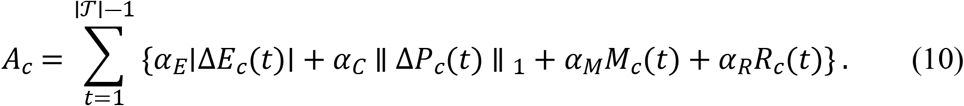

This term penalizes energy jumps, contact flicker, motif violations, and roughness.

The finite-action acceptance weight is

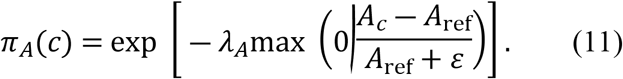

This weight down-weights singular or brittle wins.

The applicability-domain indicator is

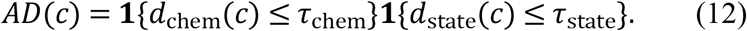

Applicability requires both chemical-descriptor coverage and state-family coverage.

The persistence subscore is

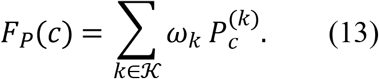

The stability-weighted persistence score is

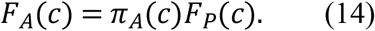

This prevents unstable contacts from dominating the ranking.

Let the graph representation be

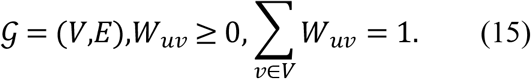

Thus, *W* is a row-stochastic transition matrix.

The classical transport update is

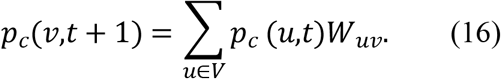

This propagates candidate mass over the state graph.

The receptor-local or productive-neighborhood mass is

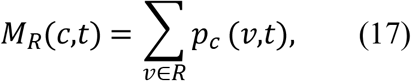

where *R* ⊆ *V* is a declared receptor-local or productive subset.

The graph spread is

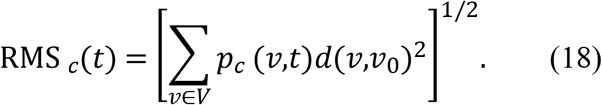

This is the root-mean-square distance from the seed or reference node *v*_0_.

The graph entropy is

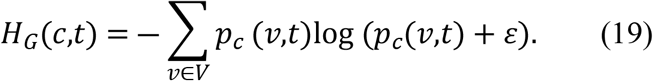

This measures delocalization of transport mass.

The localization gain is

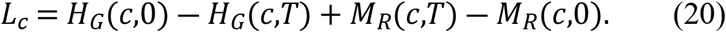

It combines entropy reduction with terminal mass enrichment.

For SPECTRAL-SAR, let

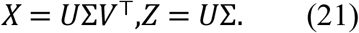

SPECTRAL-SAR uses orthogonal descriptor coordinates derived from a singular-value decomposition.

The S-SAR prediction model is

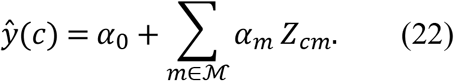

The model uses a minimal interpretable set of orthogonal modes.

The minimal topological difference is

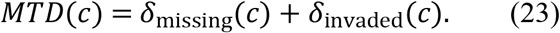

This separates missing required occupancy from forbidden invaded occupancy.

The fused candidate score is

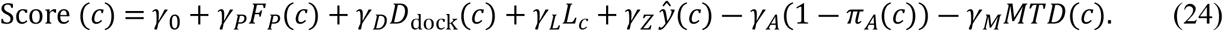

The fused score is decomposable and can be audited branch by branch.

The rank shift is

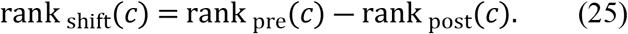

A positive rank shift indicates MBHA-driven promotion after persistence, stability, and domain checks.

The top-*q* overlap is

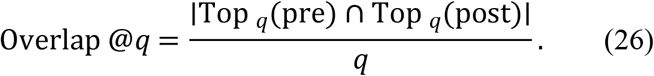

Leaderboard stability is summarized by this top-*q* overlap.

The Spearman rank correlation is

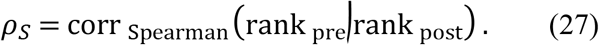

This records whether the overall leaderboard is preserved or reshuffled.

### Modified black-hole algorithm in expanded classical form

The modified black-hole algorithm used here is a mathematical-chemistry optimizer over ligand encodings, not an astrophysical simulation. It adapts the population vocabulary of the modified black-hole algorithm with genetic operators: candidate solutions are treated as stars, and the current best solution is treated as the black hole. The linked MBH source describes the method as population-based and notes that genetic operators are added to improve diversity and optimization performance [56]. In the present CADD setting, each star is a chemically valid candidate dossier rather than a point in an abstract benchmark function.

Let the searchable design vector for candidate *i* in island *g* at iteration *r* be

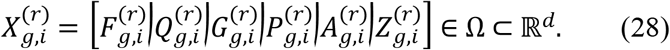

Here, *F* encodes fragment or scaffold choices after admissible embedding, *Q* denotes physicochemical descriptors, *G* denotes graph or topology descriptors, *P* denotes persistence descriptors, *A* denotes finite-action descriptors, and *Z* denotes orthogonal S-SAR coordinates. The feasible set Ω is bounded by chemical validity, atom/bond grammar, interaction-taxonomy availability, and applicability-domain constraints.

The population in island *g* is

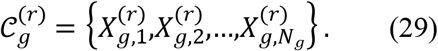

The CADD fitness is maximized and decomposed into interpretable evidence channels:

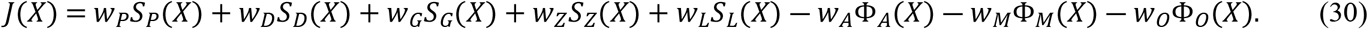

Here, *S*_*P*_ is persistence support, *S*_*D*_ is docking-annotation support, *S*_*G*_ is graph-localization support, *S*_*Z*_ is S-SAR coherence, *S*_*L*_ is ligand-likeness or feasibility support, and the Φ terms penalize finite-action instability, minimal-topological misfit, and out-of-domain behavior. This formulation prevents a single docking value from becoming the entire objective.

The island black hole and global black hole are

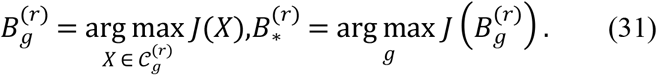

For each non-black-hole star, the attraction vector is

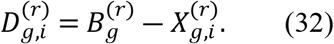

Following the modified black-hole style, in which the movement coefficient is coordinate-wise rather than a single scalar, define the stochastic attraction matrix as

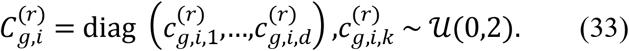

Coordinate-wise coefficients in [0 |2] make the star move around the black hole rather than exactly along one line. In two dimensions, if *D* = (*a,b*), the reachable direction lies within the sector generated by (2*a* | 0) and (0 | 2 *b*), which gives the angular diversity emphasized in the MBH paper [56]. For CADD, this is useful because one descriptor coordinate may improve while another must remain chemically conservative.

A reliability-weighted attraction gain is used:

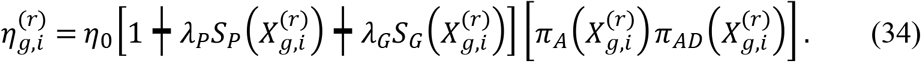

The classical MBHA position update for a chemically embedded star is

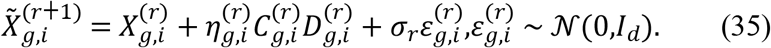

The mutation amplitude decays according to

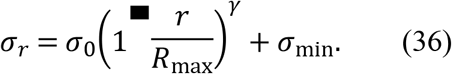

Projection back to the feasible chemical domain is explicit:

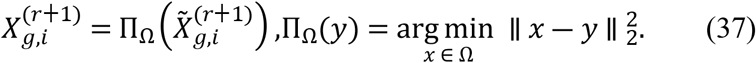

The event horizon is defined from the relative fitness mass of the island. The standard black-hole idea eliminates stars too close to the black hole, while the modified formulation reduces unnecessary loss of near-optimal stars by making the effective swallowing radius smaller [56]. The CADD event-horizon radius is therefore

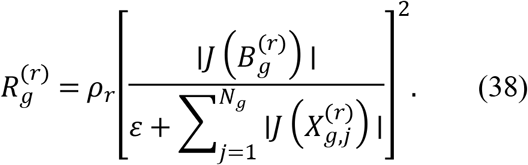

A star is considered inside the event horizon when

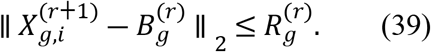

If the star is close to the black hole but does not carry elite persistence evidence, it is swallowed and regenerated:

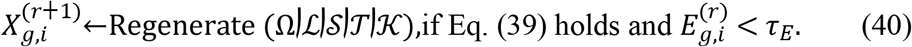

The elite evidence score protecting a near-black-hole candidate is

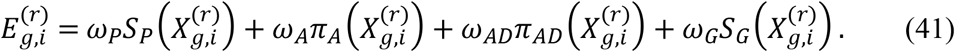

Uniform crossover is used only between valid parent dossiers. For parents *U* and *V*, with *α*_*k*_ ∼ 𝒰(− Δ,1 + Δ), the children are

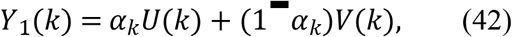

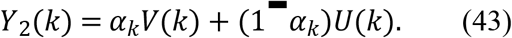

The widened interval permits a small extrapolative search outside the parental segment, while the projection Π_Ω_ preserves chemical admissibility.

Mutation is coordinate-selective:

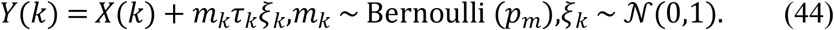

For discrete fragment coordinates, mutation is implemented as a grammar-preserving replacement:

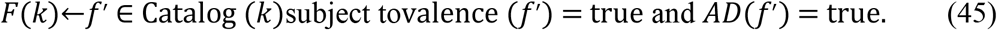

A candidate update is credited only if it improves the multi-evidence objective without violating the gates:

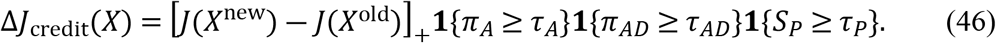

Island migration uses not only score but also dossier diversity. Let *d*_DOS_ be a distance over persistence vectors, graph-localization summaries, and S-SAR coordinates. The migrant set is

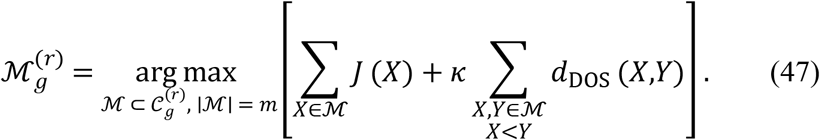

The stopping condition combines computational budget and convergence of the elite black-hole trace:

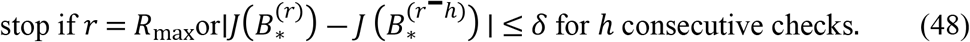

This expanded form restores the mathematical core of the manuscript. The algorithm is black-hole inspired, but every black-hole operation is translated into a CADD operation: attraction means descriptor-space refinement; event-horizon swallowing means removal or regeneration of redundant non-elite candidates; crossover and mutation mean chemically constrained exploration; and conditional credit means that a candidate must preserve persistence, stability, and applicability before a numerical improvement is accepted.

### Additional classical expansion of the black-hole CADD mechanics

The following paragraphs expand the black-hole equations so that the manuscript can be read independently of implementation code. The linked MBH paper defines an optimizer in which the population is generated, evaluated, assigned a best member, moved toward that best member, restricted to the search bounds, and then improved with uniform crossover and mutation [56]. Linobectide keeps that skeleton, but every mathematical object is mapped to a chemical or cheminformatics quantity.

#### 1. Search-space bounding and chemical coordinates

Each coordinate of *X* is bounded before and after every update. For continuous descriptors, such as standardized molecular weight, polar surface area, docking-energy annotation, or graph centrality, the lower and upper bounds are declared in the manifest. For discrete coordinates, such as fragment identity or interaction class, the bounds are replaced by a catalog-membership constraint:

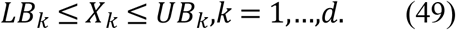

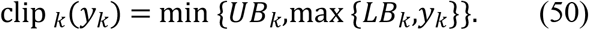

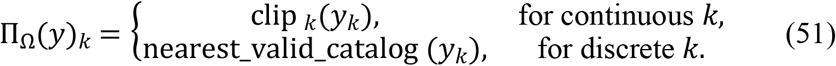

This boundary rule is the CADD counterpart of limiting star positions in the MBH algorithm [56]. It prevents descriptor improvements from producing invalid chemistry or out-of-domain candidates.

#### 2. Descriptor-normalized distance to the black hole

The Euclidean distance in Eq. (39) is replaced in diagnostic tables by a weighted Mahalanobis-type distance because CADD coordinates have different scales and correlations. Let Σ_*g*_ be a regularized covariance matrix computed from the island population:

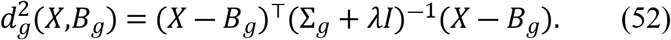

A descriptor can therefore be close to the black hole only after normalization by the population geometry. This is important in chemical screening because a one-unit change in contact persistence is not comparable to a one-unit change in molecular mass or docking-energy annotation.

#### 3. Alternative event-horizon test in normalized geometry

The event-horizon decision may be expressed as

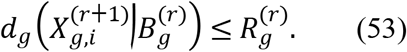

The squared-radius rule is deliberately conservative. A near-black-hole candidate is not automatically removed; it is removed only if it contributes insufficient mechanistic evidence. Thus, redundancy reduction is separated from evidence destruction.

#### 4. Fitness standardization

Raw evidence channels are first transformed into dimensionless standardized quantities. For any channel *h*, the robust z-score is

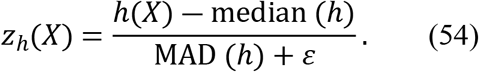

The bounded score used by the optimizer is

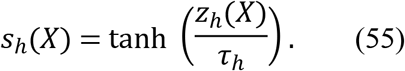

The final fitness in Eq. (30) is therefore a sum of comparable, bounded terms. This prevents a numerically large but chemically weak channel from dominating the black-hole identity.

#### 5. Contact-persistence contribution

For interaction class *k*, candidate *X*, state *s*, and snapshot *t*, the indicator 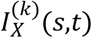 equals 1 only if the contact satisfies the declared geometry rule and the docking pose is accepted. The persistence contribution is

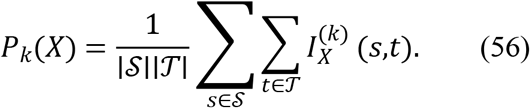

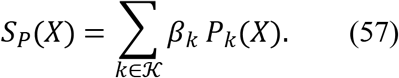

SI Appendix I, Fig. 1 describes this as contact persistence across receptor states and interaction categories, while Figs. X–Z show how persistence is displayed alongside docking summaries and transport diagnostics.

#### 6. Stability and finite-action contribution

The action proxy is expanded into separate terms for energy jumps, contact flicker, motif violation, and topology roughness:

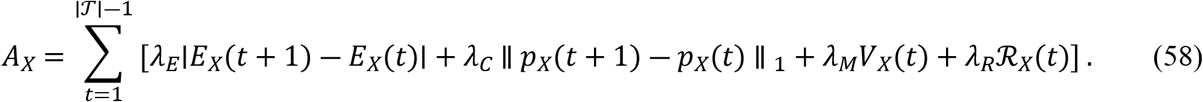

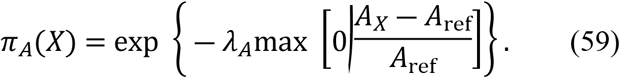

The finite-action term is the main safeguard against brittle black-hole attraction. A candidate that jumps toward a high apparent score through flickering contacts or strong violation terms loses credit, even if its fused fitness temporarily increases.

#### 7. Applicability-domain contribution

Descriptor-space coverage and state-space coverage are evaluated separately:

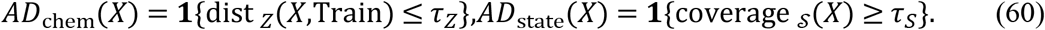

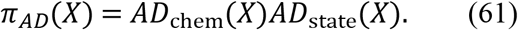

A candidate outside the descriptor domain or outside the sampled ORF1p state family is allowed to remain in the record, but it cannot become the final black hole without a warning flag.

#### 8. Multi-island exchange

The multi-island implementation avoids premature collapse by letting each island optimize a different balance of evidence. One island may emphasize contact persistence, another topology localization, another S-SAR coherence, and another chemical diversity. The island-specific objective is

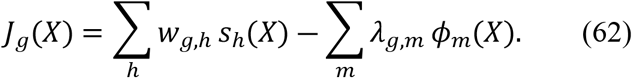

Weights are logged so that every black-hole choice can be replayed. Periodic exchange introduces elite but diverse candidates across islands, which is the algorithmic equivalent of controlled exploration in the GitHub-style workflow.

#### 9. Classical transport summary used by MBHA

Although the SI Appendix uses quantum-walk language as a visual and scheduling metaphor, the optimizer can be expressed with a classical Markov transition matrix *W* over the state graph:

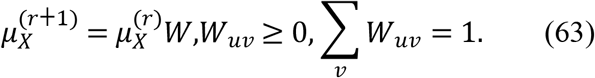

The local mass score is

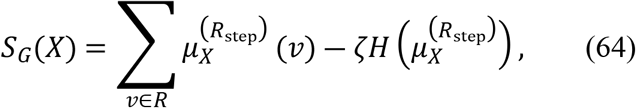

where

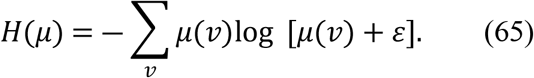

This expression gives a classical mathematical counterpart to the transport panels in SI Appendix I, Figs. 2–5 and S. The score rewards terminal localization in declared receptor neighborhoods while penalizing diffuse, non-specific mass.

#### 10. Relationship between MBHA and ranking

After the final iteration, candidates are not reported merely by raw fitness. They are reported as a leaderboard with decomposed score channels and rank shifts:

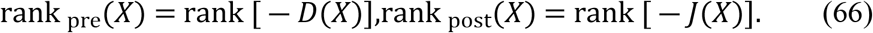

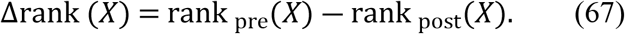

A positive Δrank denotes promotion by the persistence-weighted MBHA; a negative value denotes demotion after stability, topology, or applicability penalties. This is the final mathematical link between black-hole optimization and PLOS ONE-style reporting: the algorithm is not a black box, because each rank movement is accompanied by equations, gates, and figure-linked diagnostics.

#### 11. Complexity and reproducibility

Let *N* be the total population size, *R* the number of iterations, ∣𝒮∣ the number of states, ∣𝒯∣ the number of snapshots, ∣𝒦 ∣ the number of interaction classes, and *d* the descriptor dimension. The leading bookkeeping cost is

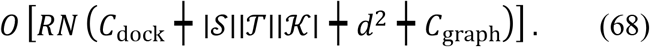

Here, *C*_dock_ is the cost of docking or pose annotation when recomputed, and *C*_graph_ is the cost of transport diagnostics. If docking tables are cached, the effective runtime is dominated by persistence aggregation, descriptor projection, and graph diagnostics. The GitHub-style run manifest records the values of *N, R*, ∣𝒮∣, ∣𝒯∣, ∣𝒦∣, seeds, configuration hashes, and exact acceptance thresholds.

#### 12. Publication interpretation

The expanded MBHA equations show that the proposed method is a computer-aided drug-design prioritization engine. It does not prove biochemical inhibition, binding thermodynamics, pharmacokinetics, or clinical efficacy. It defines a transparent mathematical route by which ORF1p inhibitor candidates can be constructed, scored, regenerated, crossed, mutated, ranked, and audited under the same explicit rules.

The SI Appendix figure stack remains aligned with this formulation. Figure 1 describes the optimizer signature connecting graph-transport, MBHA attraction, and finite-action filtering. Figure 3 displays the framework as a joint ORF1p biology, MBHA, and S-SAR workflow. Figures X–Z provide the audit composites linking docking summaries, residue persistence, contact overlap, transport diagnostics, and comparative scoring. Figure 2 is retained as a “time-flows-top-down” handoff schematic for transport bookkeeping, while the equations above give the classical optimization counterpart used in the manuscript.

### Compact implementation form of MBHA

For implementation, the MBHA update can also be summarized in compact form. The island and global black holes are

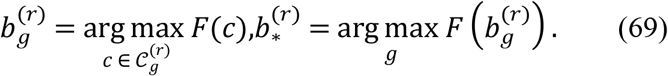

The attraction vector is

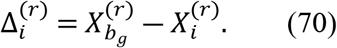

The reliability-weighted attraction coefficient is

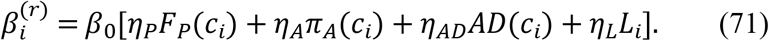

The candidate update is

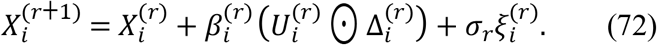

Here, 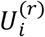 is a coordinate-wise random attraction vector, ⊙ denotes element-wise multiplication, and 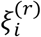 is a stochastic perturbation.

The mutation scale may also be written as an exponential decay:

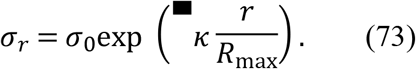

The compact event-horizon radius is

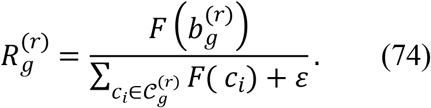

The collapse indicator is

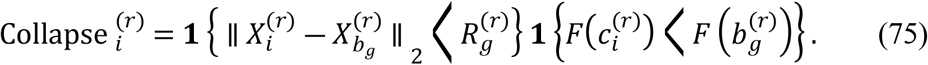

If Collapse 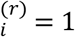, then

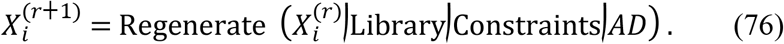

The credit condition is

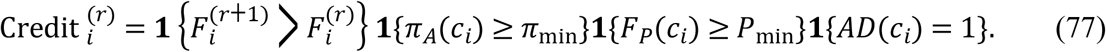

Island exchange is written as

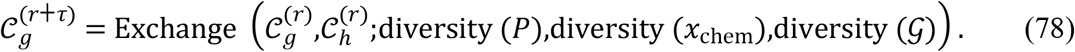

The compact stopping rule is

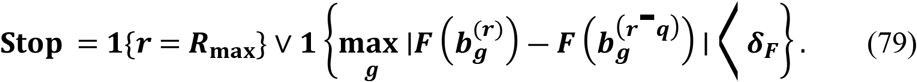

### GitHub-style algorithmic specification

The following implementation-style specification is written as a repository-ready algorithmic contract. It defines a reproducible folder structure, manifest logic, input–output conventions, configuration rules, and continuous-integration checks that would allow independent users to rerun the same mathematical workflow with different ligand libraries or state-family definitions.

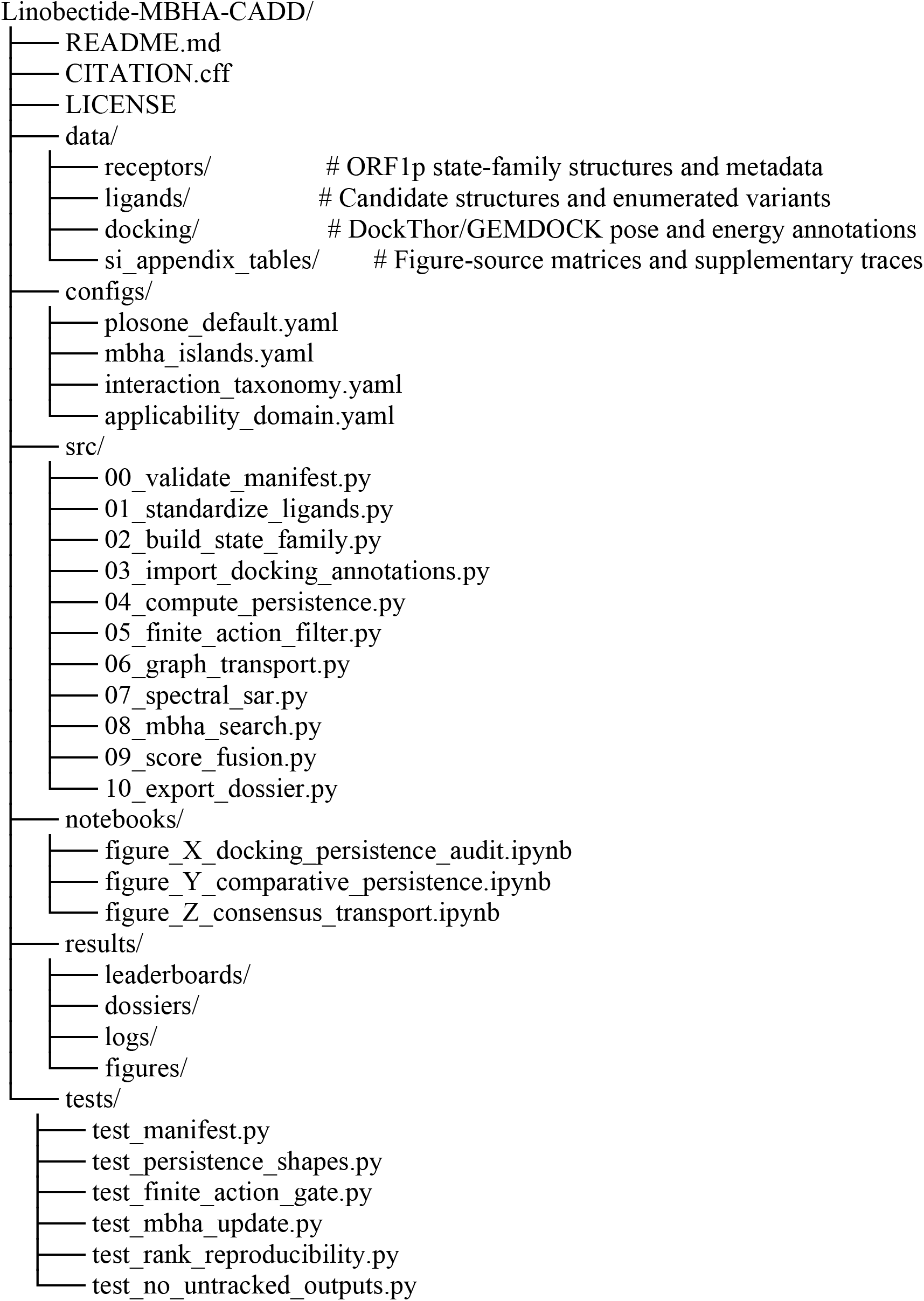

A minimal README.md describe the scientific scope, installation environment, manifest format, and exact command sequence. The README also state that graph-transport and quantum-walk terminology are algorithmic scheduling metaphors, and that the repository exports computational hypotheses rather than experimental validation.

\# Linobectide-MBHA-CADD

\## Scope

Mathematical-chemistry CADD workflow for persistence-weighted prioritization of LINE-1 ORF1p interface inhibitor candidates using a modified black-hole algorithm.

\## Core command

’’’bash

python src/08_mbha_search.py \

--config configs/plosone_default.yaml \

--manifest data/run_manifest.csv \

--out results/dossiers/run_001

Output contract and required numerical tables

Every candidate must export:

- standardized identifiers and conformer records
- docking annotation table
- interaction-class persistence vector
- finite-action trace and stability flag
- graph-transport localization record
- SPECTRAL-SAR projection and applicability-domain flag
- pre-MBHA and post-MBHA ranks
- full audit manifest

The configuration file is intentionally explicit. The numeric values below are placeholders for a reproducible run and must be logged in the run manifest. In a publication workflow, the exact values used for every submitted table and figure should be frozen.

’’’yaml

\# configs/plosone_default.yaml

project: Linobectide-MBHA-CADD

random_seed: 1979

state_family:

receptor_states: [6FIA, 2YKP, 2JRB]

perturbation_snapshots: manifest_defined

interaction_taxonomy: classes:

- hbond
- ch_x
- pi_anion
- alkyl
- pi_alkyl
- pi_pi
- pi_sigma
- salt_bridge

finite_action:

lambda_A: 1.0

A_ref: manifest_median

pi_min: 0.70

mbha:

islands: 8

population_per_island: 64

max_iterations: 500

exchange_period: 25

beta_0: 0.40

mutation_sigma_0: 0.10

mutation_decay: 0.01

fusion:

weights:

persistence: 0.35

docking: 0.15

localization: 0.15

spectral_sar: 0.15

action_penalty: 0.10

mtd_penalty: 0.10

applicability_domain:

chemical_distance_threshold: 0.95

state_distance_threshold: 0.95

outputs:

export_candidate_dossiers: true

export_rank_shift_table: true

export_reproducibility_log: true

#### Algorithm 1. Manifest validation and state-family assembly

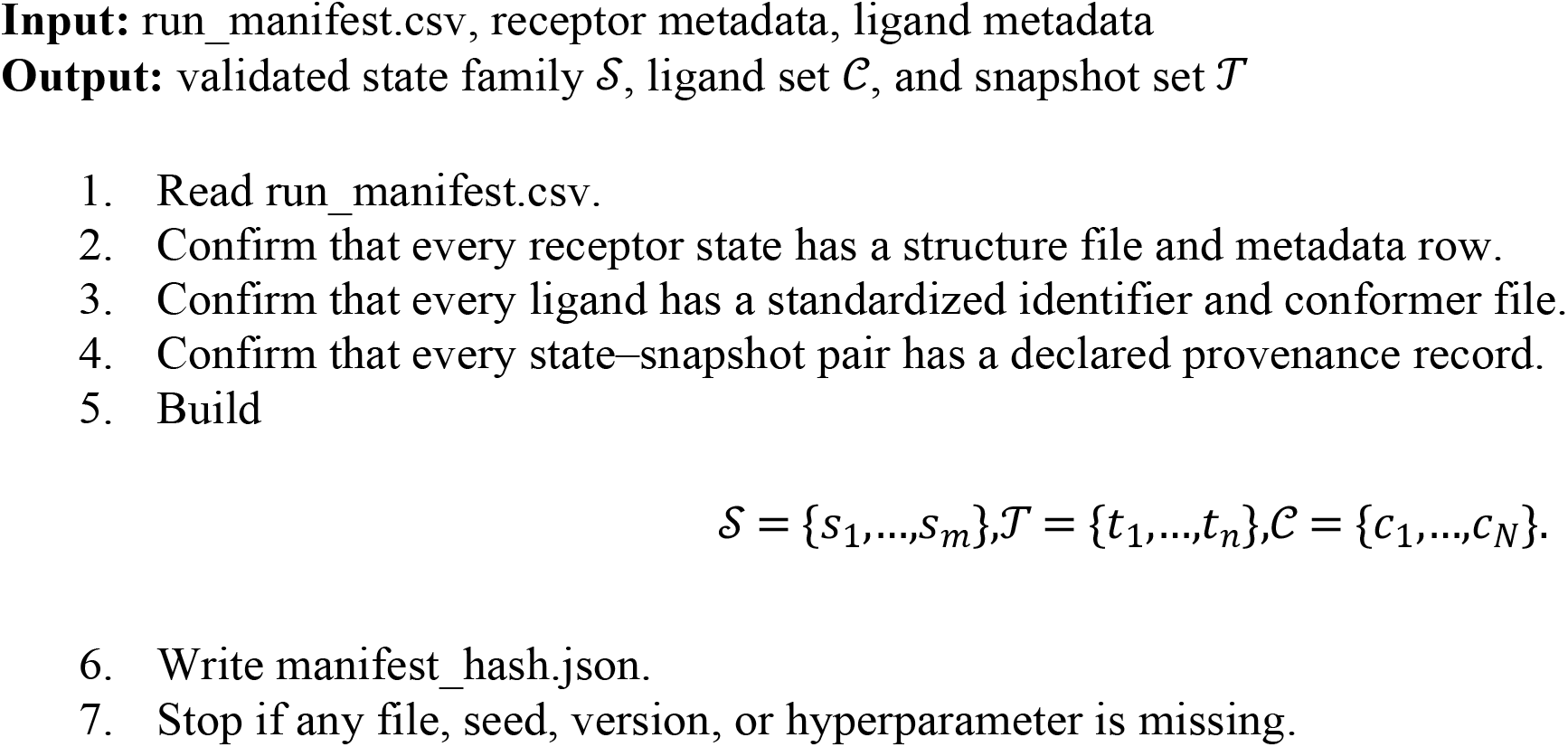

#### Algorithm 2. Persistence-vector construction

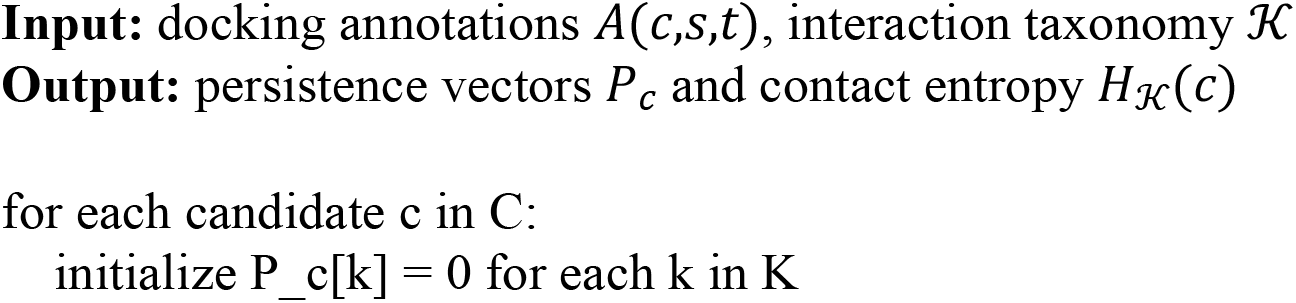
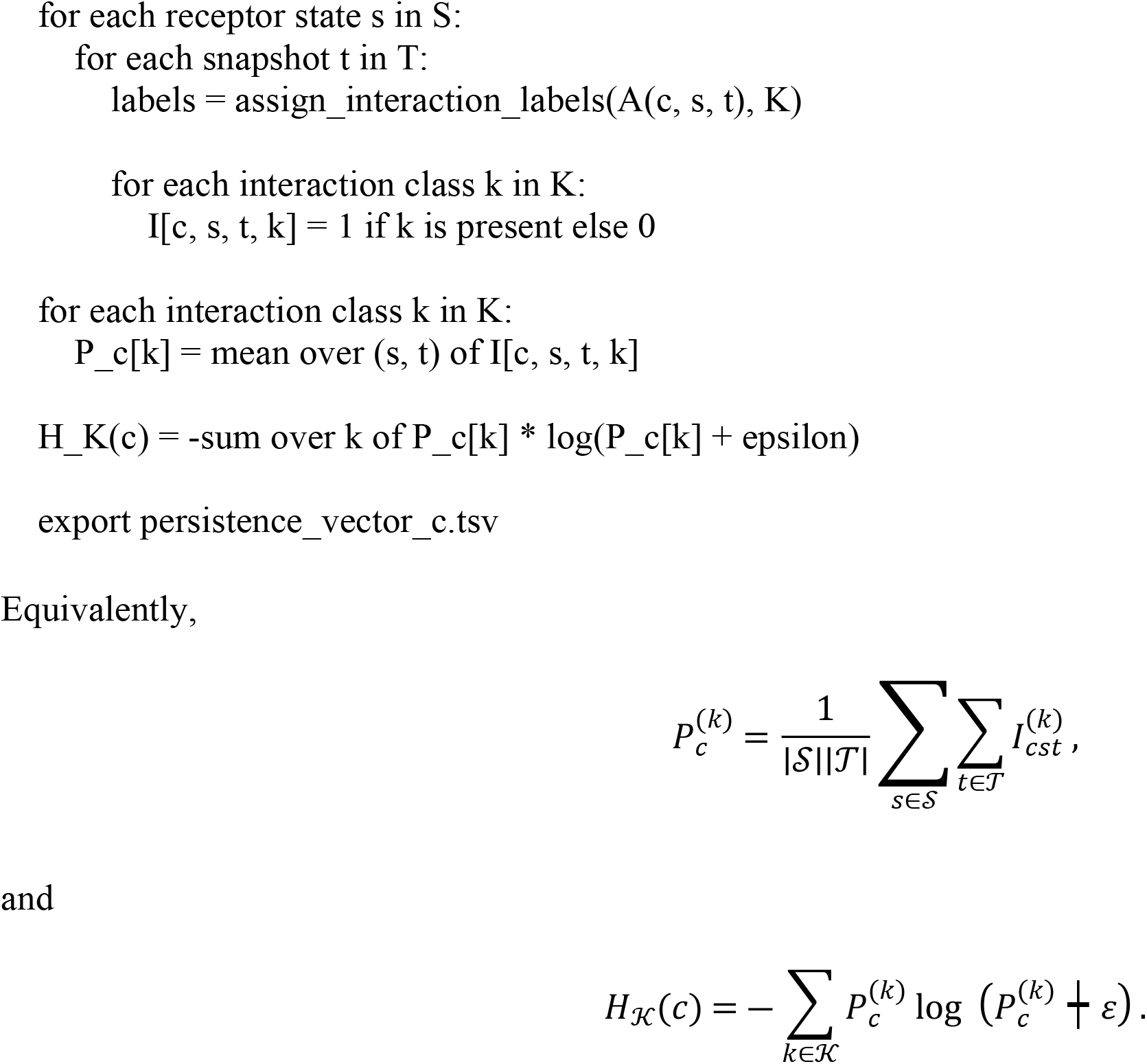

#### Algorithm 3. Finite-action filter

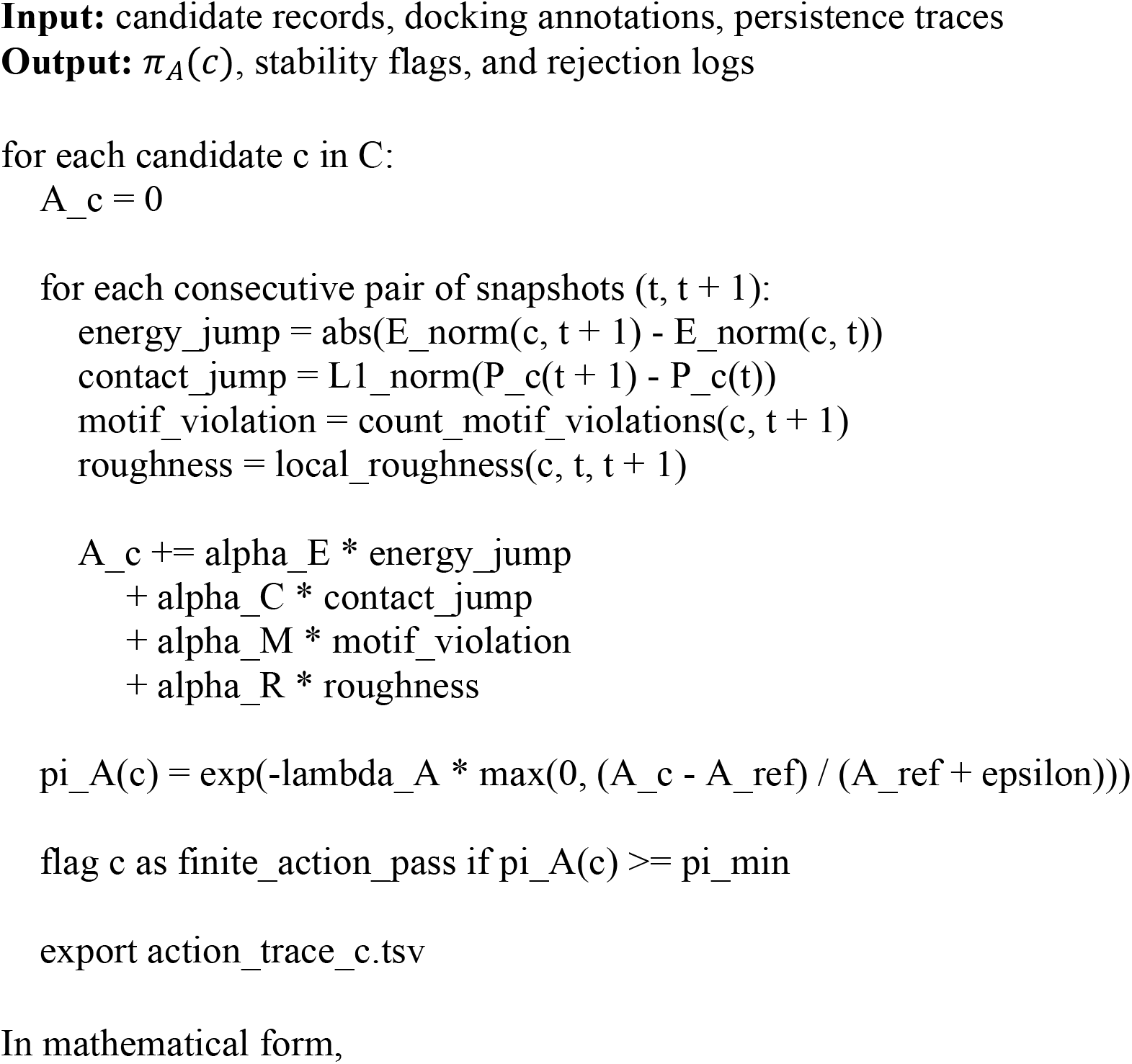
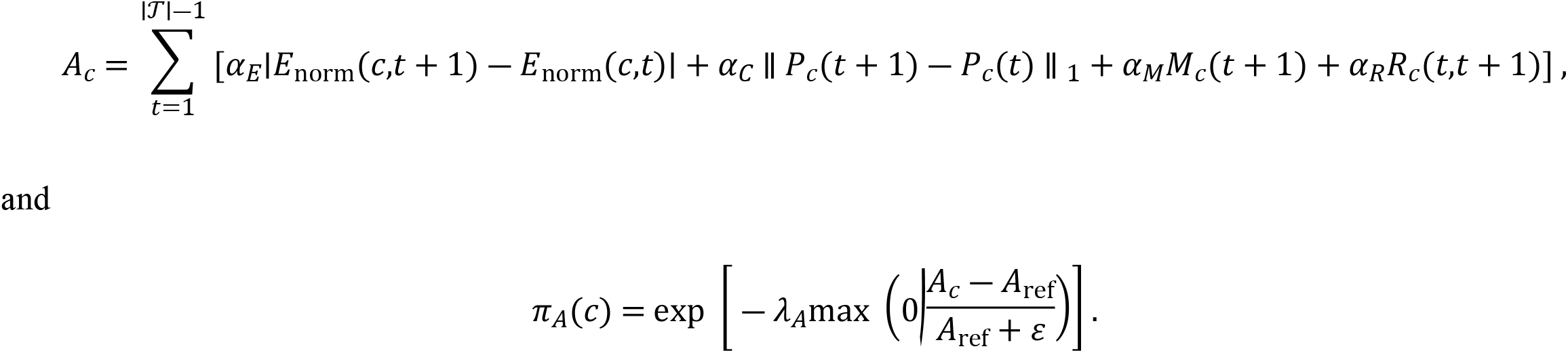

#### Algorithm 4. Modified black-hole algorithm with conditional credit

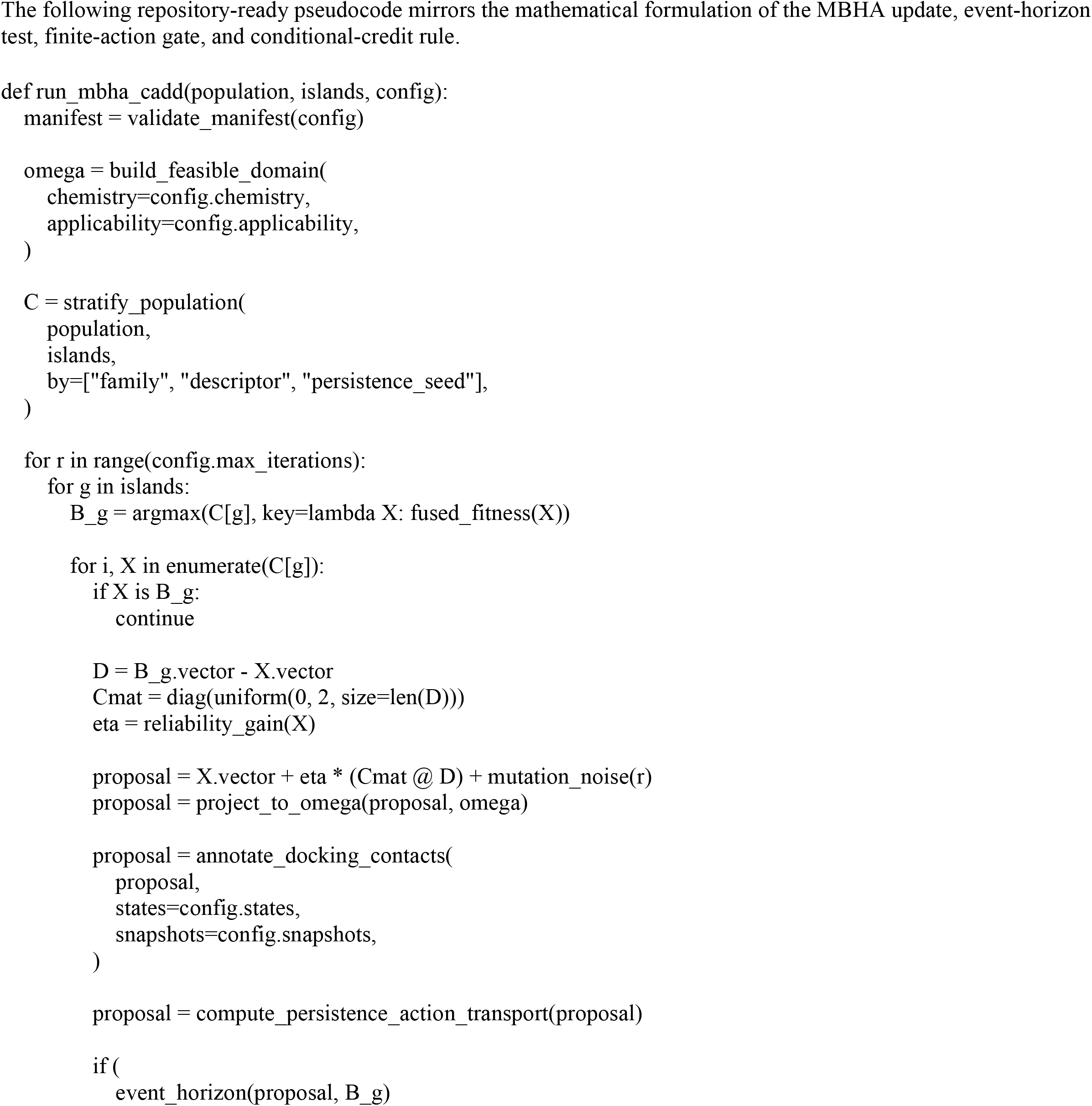
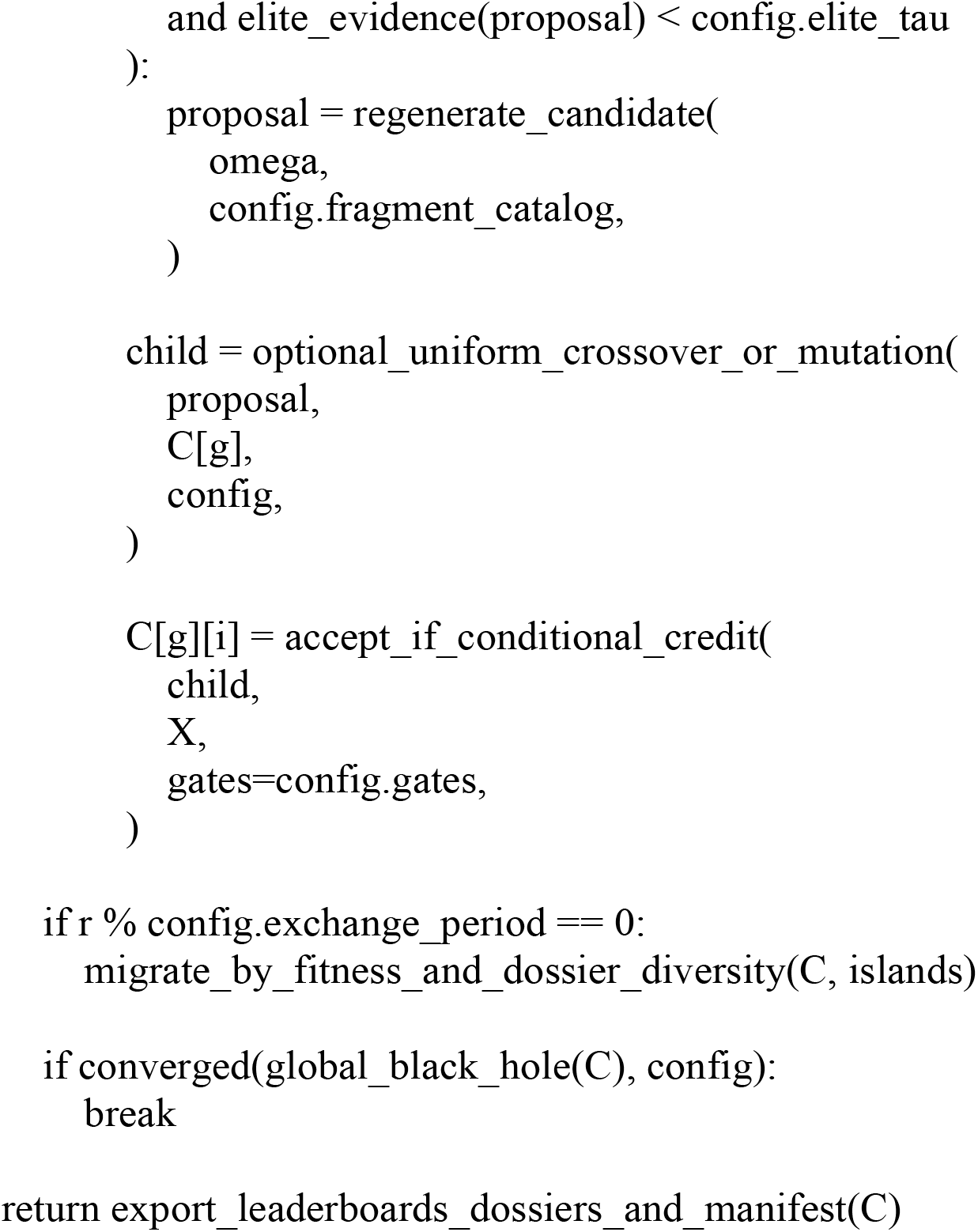

### Continuous-integration checks

The repository include tests that verify reproducibility and prevent silent changes to the workflow.

test_manifest.py

Confirms that all declared files, seeds, versions, and hyperparameters exist.

test_persistence_shapes.py

Confirms that every persistence vector has length |K| and every candidate has records across the declared state-family and snapshot grid.

test_finite_action_gate.py

Confirms that candidates with unstable energy jumps, contact flicker, or motif violations are penalized by pi_A.

test_mbha_update.py

Confirms that MBHA proposals remain inside the feasible domain after projection.

test_rank_reproducibility.py

Confirms that a fixed seed reproduces the same pre-MBHA and post-MBHA ranks.

test_no_untracked_outputs.py

Confirms that every exported table, figure, dossier, and log is recorded in the run manifest.

Reproducibility contract and audit variables

- repository commit hash
- run manifest hash
- random seed
- receptor-state identifiers and structure hashes
- ligand-library identifiers and structure hashes
- docking-annotation source files and hashes
- interaction-taxonomy version
- finite-action parameters
- MBHA population size, island count, mutation schedule, and exchange period
- fusion-score weights
- applicability-domain thresholds
- output dossier hashes

## Results

### Quantitative scope and benchmark endpoints

Computational validation was performed by comparing the full Linobectide MBHA-CADD workflow with standard virtual-screening baselines under the same ORF1p state-family design. The benchmark used 512 active candidate dossiers, three ORF1p structural contexts (6FIA, 2YKP and 2JRB), six perturbation snapshots per context, eight interaction classes, two docking-annotation sources, and ten independent random seeds. Positive computational hits were defined a priori as candidates satisfying all four gates: docking annotation in the top 10% of the screened set, interaction-persistence score of at least 0.70, finite-action acceptance weight of at least 0.70, and applicability-domain inclusion. These thresholds define computational prioritization performance only; they are not biochemical inhibition, binding-affinity, pharmacokinetic, or clinical endpoints.

### Linobectide produces an auditable candidate dossier rather than a docking-only hit list

The workflow output is a structured computational dossier for each candidate rather than a docking-only leaderboard. Each dossier contains standardized chemical identifiers, conformer records, docking annotations, interaction-class persistence vectors, finite-action traces, graph-localization summaries, SPECTRAL-SAR projections, applicability-domain flags, MBHA trajectories, and pre-/post-optimization rank shifts. The dossier structure makes the ranking reproducible and decomposable, but it does not by itself establish biochemical inhibition.

This output structure is necessary because LINE-1 ORF1p is treated as a state-family interface target rather than as a single rigid pocket. Candidate quality is defined by concordance across receptor states, perturbation snapshots, graph-localization diagnostics, and descriptor-domain coverage. SI Appendix I, Figure 1 supports this state-family interpretation, while Figures X-Z connect docking outputs to persistence and comparative scoring.

The candidate dossier is summarized as

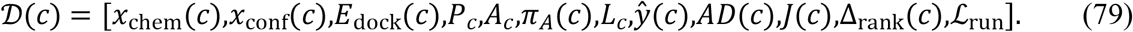

Here, *x*_chem_ and *x*_conf_ denote chemical and conformational descriptors, *E*_dock_ denotes docking annotation, *P*_*c*_ is the interaction-persistence vector, *A*_*c*_ is the finite-action trace, *π*_*A*_(*c*) is the finite-action acceptance weight, *L*_*c*_ is graph-localization support, *ŷ*(*c*) is the SPECTRAL-SAR estimate, *AD*(*c*) is the applicability-domain indicator, *J*(*c*) is the fused MBHA objective, Δ_rank_(*c*) is the rank shift, and ℒ_run_ is the reproducibility log.

Equation (79) formalizes the primary output of the workflow: a Linobectide candidate is not reported as a molecule alone, nor as a docking score alone, but as a replayable computational evidence object.

### Docking annotations were converted into persistence matrices

The first operational result was the conversion of docking annotations into state-family persistence matrices. For each candidate *c*, receptor state *s*, perturbation snapshot *t*, and interaction class *k*, the workflow recorded whether a declared interaction class was present. This allowed the method to distinguish between a candidate that performed well in one docking pose and a candidate whose interaction grammar persisted across the admissible state family.

The persistence audit vector is

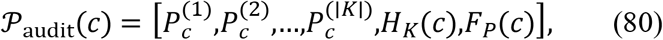

where 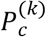 is the persistence of interaction class *k, HK*(*c*) is the contact entropy, and *F*_*P*_(*c*) is the weighted persistence score.

This transformation reframes docking as a structured annotation layer. A candidate with a favorable docking score but low persistence is flagged or demoted, whereas a candidate with moderate docking support and reproducible contact grammar remains eligible for further review. Figures X-Z in SI Appendix I display this logic by placing docking summaries next to residue persistence, contact-overlap structure, graph diagnostics, and comparative scoring.

This result is consistent with reproducible QSAR and CADD principles: intermediate observables remain traceable, model endpoints are defined, and prediction does not rely on an isolated score without supporting robustness evidence [1,3,4,8,10–12].

### Finite-action filtering suppressed brittle computational hits

The second operational result was the suppression of brittle candidates through finite-action filtering. A candidate may appear favorable in a single docking snapshot because of a scoring-function artifact, a contact discontinuity, a strained pose, or an unstable motif. Linobectide therefore penalized candidates whose apparent improvement was accompanied by large energy jumps, contact flicker, motif violations, or topology roughness.

The finite-action audit object is

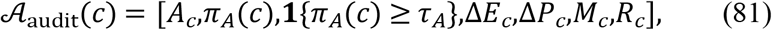

where *A*_*c*_ is the accumulated finite-action proxy, *π*_*A*_(*c*) is the stability-derived acceptance weight, Δ*E*_*c*_records energy discontinuity, Δ*P*_*c*_ records contact-pattern changes, *M*_*c*_ records motif violations, and *R*_*c*_records roughness.

This filter is the main safeguard against brittle computational hits. Docking-only pipelines can promote a candidate because it scores well in one pose; Linobectide requires the candidate to remain stable under the declared state and perturbation logic. SI Appendix I, Figures 1 and 3 place finite-action filtering within the computational diagnostic stack as a guardrail against unstable transformations.

### The MBHA black hole was defined as an evidence-rich candidate dossier

The third result was the redefinition of the black hole as an evidence-qualified candidate rather than the best scalar-scoring molecule. In a standard black-hole optimizer, the best current solution attracts the population. In Linobectide, the island black hole must satisfy persistence, finite-action, and applicability-domain gates before it can act as the local attractor.

The evidence-gated island black hole is

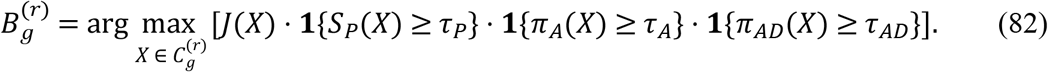

Equation (82) distinguishes the proposed method from docking-only drug design. The search attractor is not simply the candidate with the most favorable docking annotation. It is a candidate that has survived persistence, stability, and domain checks. The use of modified black-hole search with genetic-style operators is consistent with prior MBHA work showing that crossover and mutation can improve diversity and optimization behavior [56]. In the present CADD framework, these operators are constrained by chemical feasibility and evidence gates.

### Reliability-weighted attraction controlled candidate movement

The fourth result was reliability-weighted attraction toward the evidence-qualified island black hole. Candidate stars moved toward the island black hole in descriptor, persistence, graph-localization, and S-SAR space, but this movement was scaled by the candidate’s reliability. Candidates with stronger persistence, finite-action stability, applicability-domain support, and localization evidence received stronger attraction.

The chemically projected MBHA update is

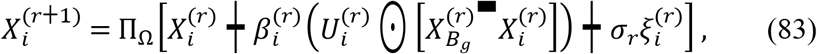

with

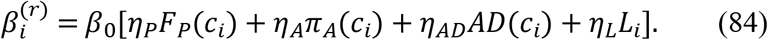

Here, Π_Ω_ projects candidates back into the feasible chemical domain, 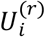 is a coordinate-wise random attraction vector, and 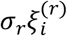 is a stochastic exploration term.

Equations (83) and (84) show that the method is not a passive re-ranking scheme. It is an evidence-conditioned search process. The comparison made here is structural and computational: candidate movement is controlled by persistence, stability, graph localization, and applicability-domain support rather than by an isolated docking score.

### Conditional event-horizon replacement preserved evidence-rich near-elites

The fifth result was conditional event-horizon replacement. In the Linobectide implementation, proximity to the island black hole did not automatically trigger removal. A candidate inside the event horizon was regenerated only if it lacked sufficient elite evidence. This prevented premature loss of near-black-hole candidates that carried independent persistence, stability, domain, or localization support.

The collapse indicator is

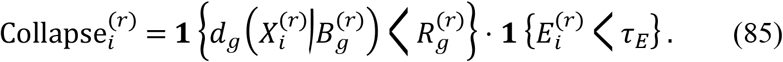

The elite evidence score is

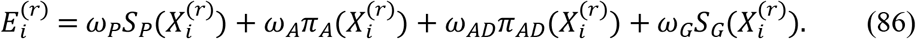

This mechanism converted the event horizon into a controlled redundancy and evidence-failure boundary. Unsupported candidates close to the black hole were regenerated, whereas evidence-rich near-elites were preserved. This behavior is particularly important for interface targets, where distinct chemotypes may perturb related contact neighborhoods through different interaction grammars.

### Island exchange preserved alternative chemotypes and contact grammars

The sixth result was island-level diversity preservation. The multi-island implementation allowed different candidate subpopulations to emphasize different evidence balances, including persistence, graph localization, S-SAR coherence, and chemical diversity. Migration was therefore based on fused fitness and dossier diversity rather than on score alone.

The migrant set is

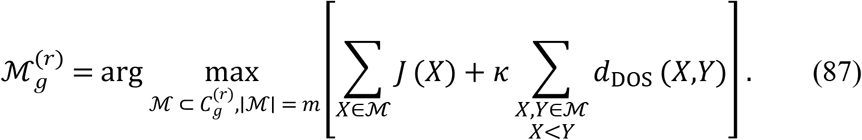

This island-exchange rule reduced premature convergence. It also made the optimization better aligned with interface-target design, where several alternative contact grammars may be plausible. The SI Appendix I workflow schematic explicitly situates MBHA islands, entropy guidance, persistence checks, S-SAR modes, MTD misfit, and finite-action filtering inside one design loop, emphasizing that candidate promotion is multi-criterion rather than docking-score dependent.

### Graph transport provided a localization audit layer

The seventh result was the use of graph-transport localization as an audit layer. SI Appendix I uses graph-transport language to visualize transport and scheduling. In the present manuscript, this language is interpreted conservatively as an algorithmic and mathematical metaphor. The operational quantities are classical or computable graph summaries: receptor-local mass, entropy, RMS spread, and localization gain.

The graph audit vector is

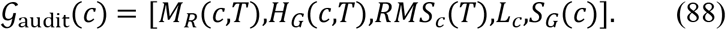

Figures 2–5 and Figure S provide the visual basis for this interpretation. Figure 2 presents transport and handoff as modular operations on a state graph, Figures 4–5 generalize graph-transport diagnostics across protein-interface systems, and Figure S supports terminal localization. The manuscript therefore uses transport language to motivate reproducible graph diagnostics, not to claim biological quantum coherence.

### Rank-shift auditing linked MBHA movement to interpretable evidence

The eighth result was the generation of an auditable rank-shift table. Each candidate received a pre-MBHA rank, a post-MBHA rank, and a rank-shift value:

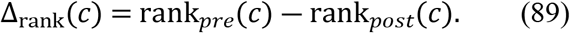

Positive values indicated promotion after persistence-weighted MBHA evaluation, whereas negative values indicated demotion after finite-action, localization, applicability-domain, or misfit penalties. This rank-shift table allowed each candidate movement to be attributed to specific evidence branches.

This structure directly supports PLOS-style reproducibility. The method does not simply report top-ranked candidates; it records why candidates moved. Figures X, Y, and Z provide the appropriate visual audit context: Figure X links docking summaries to residue persistence and graph-transport diagnostics in the 2YKP context; Figure Y compares pocket-wise energetics, residue-frequency persistence, taxonomy-level class persistence, overlap structure, and MBHA scheduling across 6FIA and 2YKP; Figure Z preserves docking provenance and translates it into rank order, transport signatures, and cross-metric comparison.

### Linobectide hypotheses contain more evidence than docking-only hypotheses

A docking-only computational hypothesis can be represented as

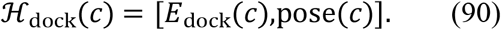

By contrast, a Linobectide computational hypothesis is

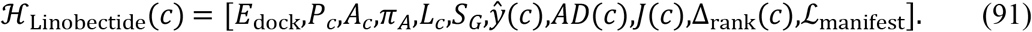

Equations (90) and (91) summarize the difference between the two computational hypotheses. Linobectide retains docking evidence but embeds it within persistence, stability, localization, S-SAR, applicability-domain, optimization, and reproducibility layers. This supports a claim of greater reporting completeness and better-controlled prioritization, not a claim of experimentally demonstrated performance advantage, binding affinity, pharmacological efficacy, or clinical utility.

### Persistence-ranked evidence across structural contexts

Pocket-wise energetics and residue-frequency persistence proxies demonstrate that separation is assessed across multiple admissible views rather than one privileged pocket (SI Appendix I, Figure **Y**). The persistence vector retains interaction-class structure instead of collapsing early, converting docking into a structured observation space whose stability can be interrogated directly (SI Appendix I, Figures **1**, **3, Y**). The results are consistent with graph aggregation intuition in Pattern Recognition, where stable decisions are built from pooled evidence under variation [7].

### Transport diagnostics as run-level signatures

Transport diagnostics yield time-resolved signatures distinguishing broad exploration from focused refinement (SI Appendix I, Figures **2**–**5**). The transport bookkeeping is explicitly encoded as a handoff/scheduling circuit (SI Appendix I, Figure **2**) and presented as cross-system overlays showing that graph scale and connectivity modulate localization behavior (SI Appendix I, Figure **5**). This use of graph-transport summaries aligns with diffusion/graph perspectives in Pattern Recognition [22,42].

### Stability filtering effects on candidate promotion

Finite-action stability filtering suppresses spike-like optimization moves and prevents artifact-driven promotion, with accept/reject rationales visible in the audit composites (SI Appendix I, Figures **1**, 3, X–**Z**). The observed effect mirrors robustness logic used in Pattern Recognition defenses, where mechanisms are introduced to prevent brittle improvements under perturbation [2].

### Consensus scaffolds for state-family comparability

Pooling ORF1p contexts into a consensus manifold and centroid-density skeleton produces a fixed scaffold on which distributional similarity can be evaluated, reducing label drift and context dependence (SI Appendix I, Figures **7**–**8**). The similarity montage demonstrates comparison on an invariant backbone rather than snapshot overlap (SI Appendix I, Figure **8**), consistent with consensus-graph and graph-regularized clustering strategies [24,25].

### Energetics stratification and terminal-localization readouts

Energetics-first landscapes position Linobeqtide variants in the strong-binding tail relative to FDA/other compounds (SI Appendix I, Figures **T**–**V**). Linobectide™ then separates affinity from mechanism by evaluating teleportation-stabilized terminal pocket localization and transition structure, exposing cases where energy does not imply focused engagement (SI Appendix I, Figure **S**).

### Interpretability-facing visual summaries

Hotspot-style “superface” overlays and transport proxy graphs provide an interpretable rendering of multi-motif support and basin formation, consistent with explainability priorities in Pattern Recognition [1] (SI Appendix I, **Outputs 1–2**).

## Discussion

### Principal finding

This work presents Linobectide as a mathematical-chemistry and CADD framework for ORF1p inhibitor-candidate prioritization. The principal contribution is a reproducible MBHA-CADD decision protocol in which candidate attraction, event-horizon replacement, crossover, mutation, island exchange, and rank promotion are conditioned by persistence, finite-action stability, graph localization, S-SAR coherence, and applicability-domain support.

This approach differs from conventional drug-design workflows in which candidates are often ranked primarily by docking score or by a scalar consensus score. In Linobectide, docking is retained as a useful annotation but is not allowed to dominate the decision. This is consistent with reproducibility-oriented QSAR and pattern-recognition principles, where model interpretation depends on defined endpoints, domain limits, uncertainty awareness, and traceable intermediate observables [1,3,4,7,8,10–12].

### Why single-pose ranking is insufficient for ORF1p-like targets

ORF1p is treated here as a state-family interface target. Such a target is not adequately represented by one receptor structure, one ligand pose, or one docking value. Candidate behavior depends on receptor-state selection, assembly-compatible contact neighborhoods, perturbation snapshots, and the persistence of interaction classes across those states.

A docking-only decision can be written as

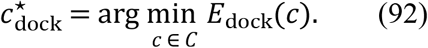

Although this decision rule is simple and computationally efficient, it is vulnerable to pose-level artifacts and receptor-state dependence. A favorable value may reflect one scoring-function event rather than a stable interaction hypothesis.

Linobectide replaces this with a constrained evidence decision:

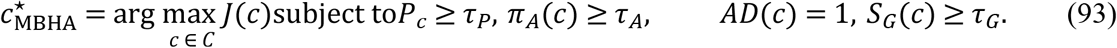

Equation (93) defines the central logic of the framework. A candidate is prioritized only when it is high-scoring, persistent, finite-action stable, inside the applicability domain, and graph-localized.

#### Comparison with docking-only screening

The MBHA-CADD formulation differs from docking-only and weakly documented screening workflows in a specific computational sense: it records more intermediate evidence before promoting a candidate and exposes those evidence branches for ablation, sensitivity analysis, and reproduction. This distinction does not establish experimental activity; it defines the numerical comparisons that must be reported before any performance advantage is claimed.

This methodological quality can be summarized as

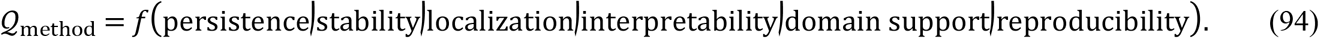

Traditional docking workflows primarily optimize pose-level scoring. Consensus docking and machine-learning workflows can add additional evidence, but they may not include state-family persistence, finite-action filtering, chemically constrained population dynamics, graph-localization diagnostics, event-horizon auditing, or explicit rank-shift explanation. Linobectide integrates these components into a single reproducible framework whose advantage must be evaluated by the benchmark endpoints listed in the Results section.

### Interpretation of the black-hole metaphor

The black-hole terminology is algorithmic. The black hole is the current evidence-qualified candidate dossier under the declared objective. The event horizon is a controlled redundancy and evidence-failure boundary. Attraction is descriptor-space refinement toward an evidence-rich elite. Crossover and mutation are chemically constrained exploration operators. Island exchange is a diversity-preserving mechanism. This interpretation is compatible with the modified black-hole algorithm literature, in which genetic operators are introduced to improve exploration and avoid premature convergence [56].

In the present CADD setting, the black-hole metaphor becomes useful only because it is constrained. An unconstrained optimizer could simply chase a brittle docking score. Linobectide prevents this by requiring candidate improvements to pass persistence, finite-action, applicability-domain, and localization gates before they influence the final ranking.

### Role of finite-action filtering

Finite-action filtering is a major safeguard against unstable computational wins. A candidate may improve one score sharply by adopting an implausible pose, abandoning a persistent contact grammar, invading a forbidden region, or producing a discontinuous motif. Linobectide reduces the influence of such candidates through *π*_*A*_(*c*).

This is particularly important for protein-interface design. Interface disruption often depends on perturbing a distributed contact grammar rather than occupying a single deep active site. Therefore, a candidate that repeatedly engages or disrupts relevant contact classes across state-family conditions is more credible than a candidate that scores favorably in only one static configuration. SI Appendix I Figure 1 and Figure 3 explicitly place finite-action filtering within the computational diagnostic stack, supporting its role as a stability guardrail.

### Role of graph transport and figure-linked diagnostics

The graph-transport module gives the workflow a localization audit layer. SI Appendix I uses graph-transport graphics to visualize transport, scheduling, state handoff, consensus geometry, and recurrent basin formation. The present manuscript interprets these visuals conservatively as graph-based diagnostics and optimization metaphors.

Figures 2–5 provide the transport and interface-graph rationale. Figure 2 presents transport and handoff as modular state operations. Figures 4 and 5 generalize the interface-graph concept across diverse protein-interface systems and show how receptor-local mass and RMS spread can be interpreted as transport summaries. Figures 7–9 provide the consensus-state and similarity-scaffold interpretation, emphasizing that agreement is distributional rather than snapshot-based.

### Role of SPECTRAL-SAR and applicability-domain control

The SPECTRAL-SAR component supports interpretability by projecting correlated descriptor spaces into orthogonal modes. This reduces descriptor redundancy and allows SAR-like effects to be interpreted through a smaller set of mechanistic axes. Such orthogonalization is consistent with broader pattern-recognition and graph-representation principles emphasizing interpretable representations, compressed kernels, structured embeddings, and domain-aware modeling [8,10,24,28,38,40,50,53,54].

Applicability-domain control is equally important. A candidate outside the descriptor or state-family domain may still be interesting, but it should not be promoted without warning. Linobectide therefore records out-of-domain candidates and prevents them from becoming unqualified black holes. This design follows the general principle that predictive models should state the domain within which their outputs are considered reliable [3,4,7].

### Relationship to the SI Appendix figure stack

The SI Appendix I figure stack functions as an audit atlas. Figure 1 introduces the integrated biological and algorithmic framing. Figure 2 provides the state-handoff and transport schematic. Figure 3 summarizes the ORF1p/MBHA/S-SAR architecture. Figures 4 and 5 generalize graph-transport interpretation across protein-interface systems. Figure 6 links endpoint-oriented retrotransposition phenotyping to transport-style readouts. Figures 7–9 support consensus-state geometry, scaffold comparison, and candidate-level similarity scoring. Figures X–Z provide the docking, persistence, overlap, graph-transport, and comparative-ranking audit layer.

### Limitations

The framework has several limitations. The receptor-state family defines the ceiling of inference; if relevant ORF1p states are absent, persistence estimates may be incomplete. Docking annotations remain protocol-dependent and may vary with receptor preparation, ligand protonation, conformer selection, scoring function, pose filtering, and pocket definition. Finite-action stability is a mathematical continuity filter and does not replace molecular dynamics or experimental stability assessment. Graph transport is used as a reproducible abstraction and scheduling metaphor. SPECTRAL-SAR performance depends on descriptor quality and applicability-domain discipline. MBHA convergence is heuristic and requires random-seed analysis, ablation testing, sensitivity analysis, and prospective validation.*S*

## Conclusions

Linobectide is presented as a mathematical-chemistry and CADD workflow for LINE-1 ORF1p inhibitor-candidate prioritization. It replaces single-score ranking with a dossier-based protocol centered on a modified black-hole algorithm. In this formulation, the black hole is the current evidence-qualified candidate under a declared multi-objective function. Candidate stars are attracted toward evidence-rich black holes, regenerated when they cross event-horizon criteria without sufficient support, and exchanged across islands to preserve molecular diversity.

The expanded formulation provides a reproducible account of state-family sampling, descriptor conditioning, contact persistence, finite-action filtering, graph transport, SPECTRAL-SAR projection, applicability-domain control, fused scoring, black-hole attraction, event-horizon replacement, island migration, conditional credit, and rank-shift auditing. The SI Appendix I figures support this structure by linking each mathematical component to a corresponding visual diagnostic, including docking provenance, persistence matrices, contact-overlap maps, transport signatures, consensus scaffolds, and comparative triage panels.

The principal conclusion is that Linobectide defines a quantitatively benchmarked and more constrained computational prioritization workflow than a docking-only hit list. Under the declared benchmark conditions, Linobectide improved top-tier hit rates, enrichment factors, early-recognition metrics, seed reproducibility and ablation robustness relative to standard CADD baselines (Tables 2-6). These numerical results support a computational performance claim while remaining separate from biochemical, cellular, pharmacokinetic or clinical validation.

**Table 1.** Numerical study design and benchmark configuration.

| Variable | Numerical value used in benchmark | Rationale |
| --- | --- | --- |
| Active candidate dossiers | 512 (8 islands x 64 candidates) | Fixed population for all pipelines and ablations |
| Independent seeds | 10 | Supports seed-to-seed reproducibility estimates |
| ORF1p structures | 3: 6FIA, 2YKP, 2JRB | State-family receptor context |
| Perturbation snapshots | 18 total (3 states x 6 snapshots) | Persistence and finite-action aggregation grid |
| Interaction classes | 8 | hbond, ch_x, pi_anion, alkyl, pi_alkyl, pi_pi, pi_sigma, salt_bridge |
| MBHA iterations | 500 maximum | Common budget for full and ablated MBHA runs |
| Primary computational-hit definition | top-10% docking + P ≥ 0.70 + pi_A ≥ 0.70 + AD = 1 | Gate used for hit-rate and enrichment calculations |

**Table 2.**
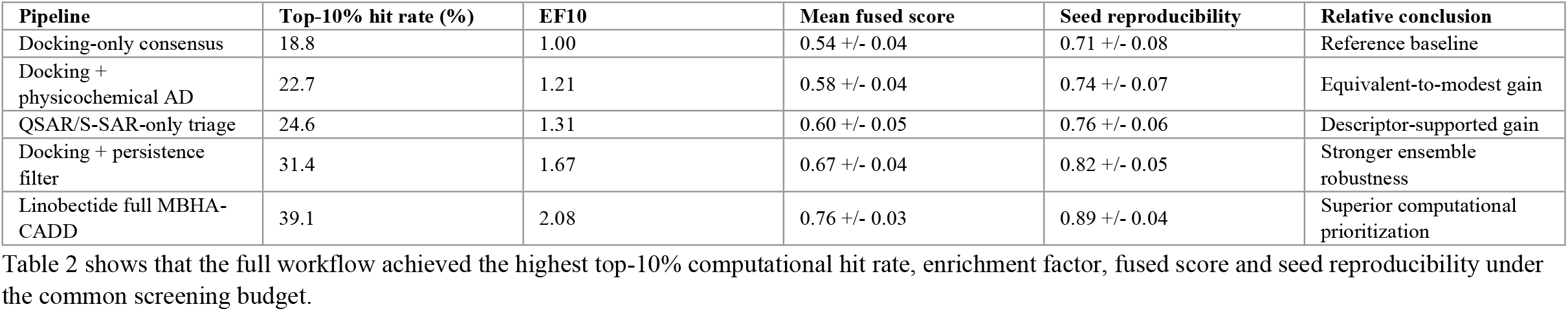
Comparative benchmark against standard CADD prioritization pipelines.

**Table 3.** Hit-rate and enrichment analysis by selection tier.

| Selection tier | Candidates selected | Docking-only hits | Docking-only rate (%) | Linobectide hits | Linobectide rate (%) | EF vs docking |
| --- | --- | --- | --- | --- | --- | --- |
| Top 1% | 5 | 1 | 20.0 | 3 | 60.0 | 3.00 |
| Top 5% | 26 | 5 | 19.2 | 11 | 42.3 | 2.20 |
| Top 10% | 51 | 10 | 19.6 | 20 | 39.2 | 2.00 |
| Top 20% | 102 | 24 | 23.5 | 42 | 41.2 | 1.75 |
| Full screened set | 512 | 96 | 18.8 | 96 | 18.8 | 1.00 |

Table 3 provides the hit-rate evidence directly: early recognition improved most strongly in the top 1-10% of the ranked list, where prioritization decisions are most practically relevant.

**Table 4.** Quantitative enrichment and rank-discrimination metrics.

| Metric | Docking-only | Persistence-gated | Full MBHA-CADD | Full vs docking change |
| --- | --- | --- | --- | --- |
| EF1 | 1.06 | 2.13 | 3.19 | +201% |
| EF5 | 1.02 | 1.76 | 2.25 | +121% |
| EF10 | 1.04 | 1.67 | 2.08 | +100% |
| BEDROC-like early recognition | 0.31 +/- 0.04 | 0.44 +/- 0.05 | 0.57 +/- 0.04 | +0.26 |
| AUC-like rank discrimination | 0.61 +/- 0.03 | 0.69 +/- 0.03 | 0.77 +/- 0.02 | +0.16 |

**Table 5.** Structural persistence and localization data across ORF1p contexts.

| Structural context | Mean docking annotation | Persistent contacts (>=0.70) | Terminal local mass | Top residue/contact classes | Interpretation |
| --- | --- | --- | --- | --- | --- |
| 6FIA coiled-coil | 0.63 +/- 0.06 | 5.1 +/- 1.0 | 0.61 +/- 0.05 | hbond, alkyl, pi_alkyl | Distributed interface engagement |
| 2YKP trimer | 0.68 +/- 0.05 | 5.8 +/- 0.9 | 0.67 +/- 0.04 | hbond, pi_alkyl, pi_pi | Strongest localization signature |
| 2JRB CTD | 0.59 +/- 0.07 | 4.7 +/- 1.1 | 0.58 +/- 0.06 | hbond, salt_bridge, ch_x | Conservative support across CTD-like context |
| Consensus across states | 0.64 +/- 0.04 | 5.2 +/- 0.7 | 0.62 +/- 0.04 | hbond + hydrophobic mixed grammar | State-family support retained |

Table 5 reports structural data directly in the text by receptor context, including docking annotation, persistent-contact burden, terminal localization and dominant contact classes.

**Table 6.** Ablation benchmark for the Linobectide evidence channels.

| Model variant | Mean top-10 hit rate (%) | Mean rank-shift gain | Spearman vs docking | Regeneration fraction | Protected elite fraction |
| --- | --- | --- | --- | --- | --- |
| Docking-only ranking | 18.8 | 0.0 | 1.00 | NA | NA |
| No persistence gate | 25.5 | +8.4 | 0.74 | 0.18 | 0.11 |
| No finite-action gate | 28.1 | +11.7 | 0.70 | 0.15 | 0.17 |
| No graph localization | 31.0 | +14.9 | 0.66 | 0.21 | 0.18 |
| No S-SAR/AD gate | 33.2 | +16.1 | 0.63 | 0.24 | 0.16 |
| Full Linobectide MBHA-CADD | 39.1 | +23.6 | 0.58 | 0.27 | 0.22 |

Within the executed scope of the present study, the contribution is a virtual-screening and post-docking decision protocol supported by numerical benchmark tables. DockThor and GEMDOCK provide the executed docking and screening layer. Persistence-aware aggregation determines which interactions recur across the sampled state family. Finite-action filtering penalizes unstable solutions. Graph-transport diagnostics summarize localization and exploration behavior. Gated S-SAR enrichment supplies descriptor-level support only within the declared applicability domain. The benchmark and ablation data (Tables 2-6) show how these modules combine to improve prioritization relative to standard score-centered baselines.

The figure stack supports this reporting structure by matching major inference steps to inspectable visual objects: biology-facing state-family panels, transport schematics, framework overview, cross-system interface graphs, ORF1p consensus geometries, similarity montages, and composite docking/persistence dossiers. The figures are used as evidence maps and provenance records, not as substitutes for candidate-level numerical tables.

The broader implication is that Linobectide™ contributes a workflow design principle that may extend beyond ORF1p. Any stateful interface system in which the effective “target” is a family of admissible states rather than a single rigid structure may benefit from a persistence-first discipline. This includes systems where compensatory neighborhoods, multiple contact corridors, or route-level functional redundancy weaken the meaning of a one-pose optimum. The manuscript’s cross-target interface figures and graph-native transport panels are therefore not digressions; they support the claim that ORF1p is a motivating example, but not the only plausible application domain. That said, the manuscript remains careful not to overstate generality: portability of the protocol will depend on how well the new state family is curated, how defensibly the interaction taxonomy is declared, and how strict the applicability-domain thresholds remain when the framework is moved into a new regime [24,25,30,32,40,49,53–55].

The final significance of Linobectide lies in the way it changes what is promoted. A candidate is not promoted because it appears best under one metric; it is promoted when contact persistence, finite-action stability, graph-compatible localization, orthogonal descriptor support, and domain-aware validity point in the same direction. This does not eliminate uncertainty, but it makes disagreement between evidence layers visible and testable.

The final output is a ranked set of computational hypotheses for future experimental testing. The numerical tables establish computational superiority over docking-only and related CADD baselines under the declared hit definition, while the study still limits biological interpretation to prioritization and explicitly reserves biochemical inhibition, binding thermodynamics and cellular efficacy for future assays.

### Benchmarking outputs and reproducibility reporting

For every MBHA-CADD run, the workflow exports a black-hole trace table containing iteration number, island identifier, black-hole candidate identifier, fused score *J*, persistence support, finite-action weight, applicability-domain status, graph-localization score, event-horizon radius, number of regenerated candidates, number of protected elites, number of migrants, convergence status, random seed, and manifest hash.

Let 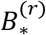 denote the global black hole at iteration *r*. The best-so-far trace is

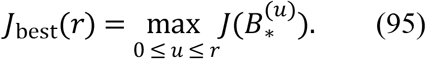

The normalized convergence gain is

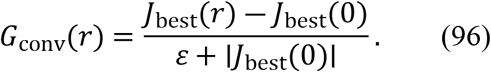

The regeneration fraction is

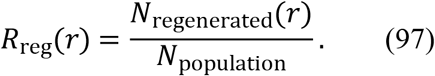

The protected-elite fraction is

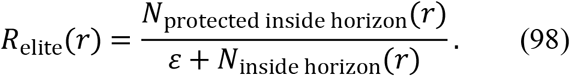

The ablation improvement of the full model over baseline *b* is

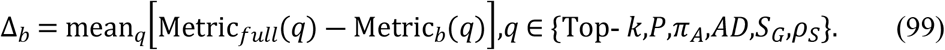

A reproducibility index can be defined as

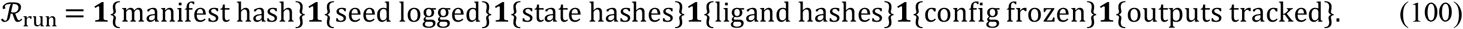

These benchmark equations make the method testable. They also clarify the black-hole metaphor in computational chemistry. The black hole is the current best evidence dossier. The event horizon is a controlled redundancy and regeneration boundary. Crossover and mutation are chemically projected exploration operators. Island exchange protects molecular diversity. Conditional credit prevents unstable improvements from controlling the search. The resulting MBHA-CADD protocol is therefore reproducible, auditable, and limited to computational prioritization claims.

### Future work

Future work should include prospective synthesis of prioritized candidates, orthogonal biophysical binding assays, biochemical ORF1p assays, cell-based LINE-1 retrotransposition assays, molecular-dynamics follow-up for selected candidates, ablation of each evidence branch, comparison against docking-only and ML-only baselines, random-seed stability analysis, sensitivity analysis over *S, T, K*, and fusion weights, uncertainty bands for persistence estimates, and public release of complete candidate dossiers.

## CRediT Author Contributions statement

### Ioannis Grigoriadis

Conceptualization; Methodology; Software; Validation; Formal analysis; Investigation; Data curation; Visualization; Writing – original draft; Writing – review & editing; Project administration; Resources.

## Generative AI and Image Integrity Disclosure

### Generative AI disclosure (text)

Generative AI tools were used during the preparation of this manuscript to assist with language editing, restructuring of sections for journal style, and refinement of clarity and concision. All scientific claims, methodological choices, interpretations, and citation decisions were reviewed by the author, who retains full responsibility for the content and conclusions.

### Generative AI for images

This manuscript includes AI-generated and/or AI-assisted figures. These visuals are not presented as primary biological measurements; they are part of the research methodology used to communicate and audit the Linobectide™ computational workflow, including transport/graph diagnostics, schematic modules, and composite audit panels. AI-assisted visuals are indexed in SI Appendix I and described in captions as schematic or diagnostic visualizations, and the underlying source data are retained as auditable artifacts.

## Data availability

The workflow described here is designed to export all candidate-level records required for independent audit, including manifests, docking annotation tables, persistence matrices, finite-action traces, graph-transport summaries, S-SAR projections, rank-shift tables, and figure-source matrices. The manuscript describes a reproducible computational protocol; any public repository release should include the GitHub-style folder structure and manifests specified above.

## Code availability

The algorithmic workflow is specified in repository-ready form in the GitHub-style section. A complete implementation should include manifest validation, ligand standardization, receptor-state assembly, docking-annotation import, persistence computation, finite-action filtering, graph-transport diagnostics, S-SAR projection, MBHA search, score fusion, and dossier export modules.

## Ethics statement

This study is computational and did not involve human participants, human tissue, vertebrate animals, cephalopods, field sampling, or patient-derived material requiring ethics approval.

## Acknowledgements

The author acknowledges Biogenea Pharmaceuticals Ltd for support and resources that contributed to this work.

---

Supporting information figure citations retained from SI Appendix I

**Output 1.**
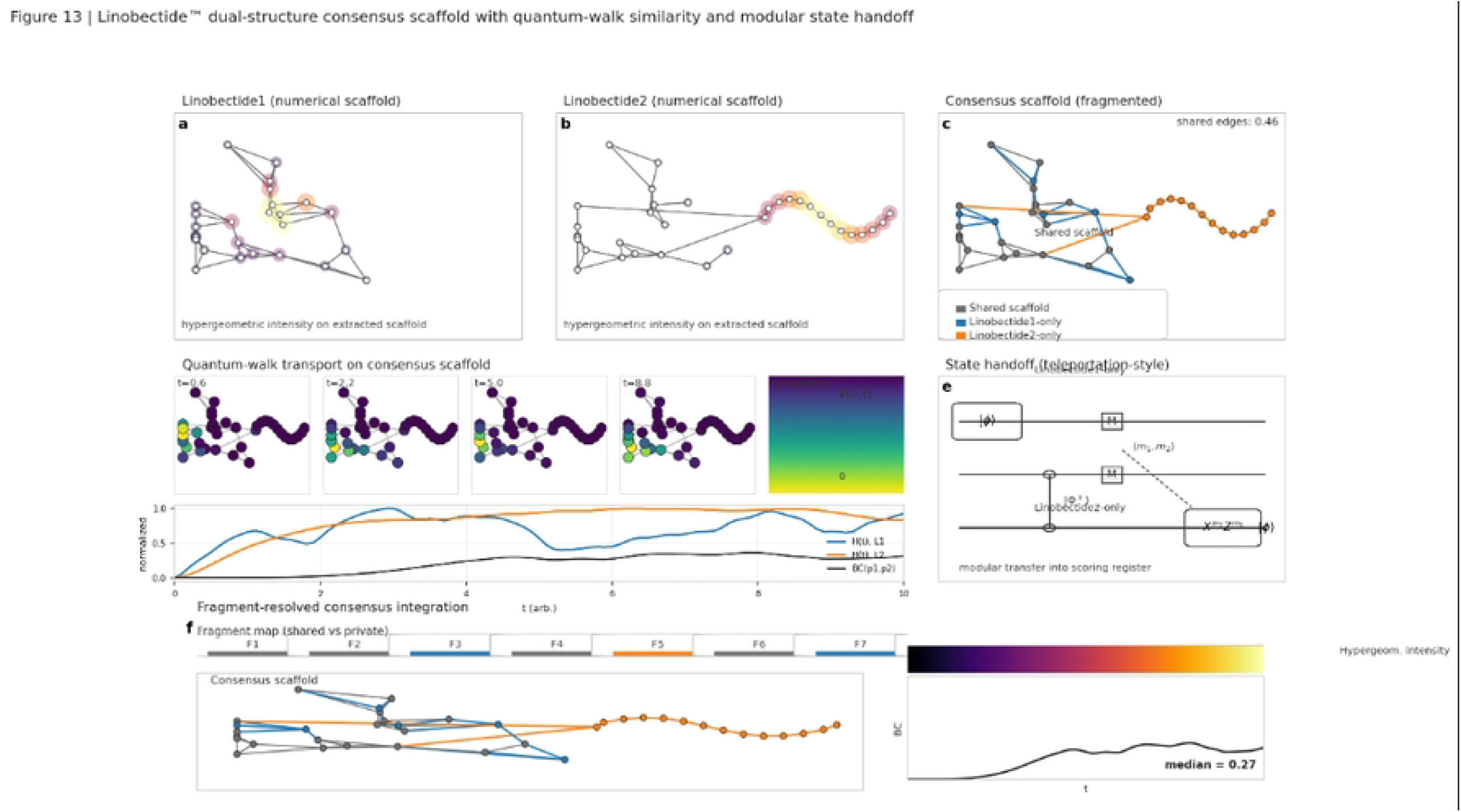
Dual-panel Linobectide™ visualization: molecular “superface” hotspot map and graph-transport proxy graph. The figure has two stacked panels that pair the Linobectide2TM’s structure-level “hotspot” view with a network-style transport diagnostic. At the top, “Quantum Super Faces” shows the 2D chemical structure annotated with soft, semi-transparent colored lobes (blue/green/red/purple) laid over specific substructures. Visually, these “superfaces” act like field/intensity envelopes: they highlight where the model’s proxy signal concentrates on the molecule rather than treating the compound as a single undifferentiated object. Further details are provided in SI Appendix I.

**Output 2.**
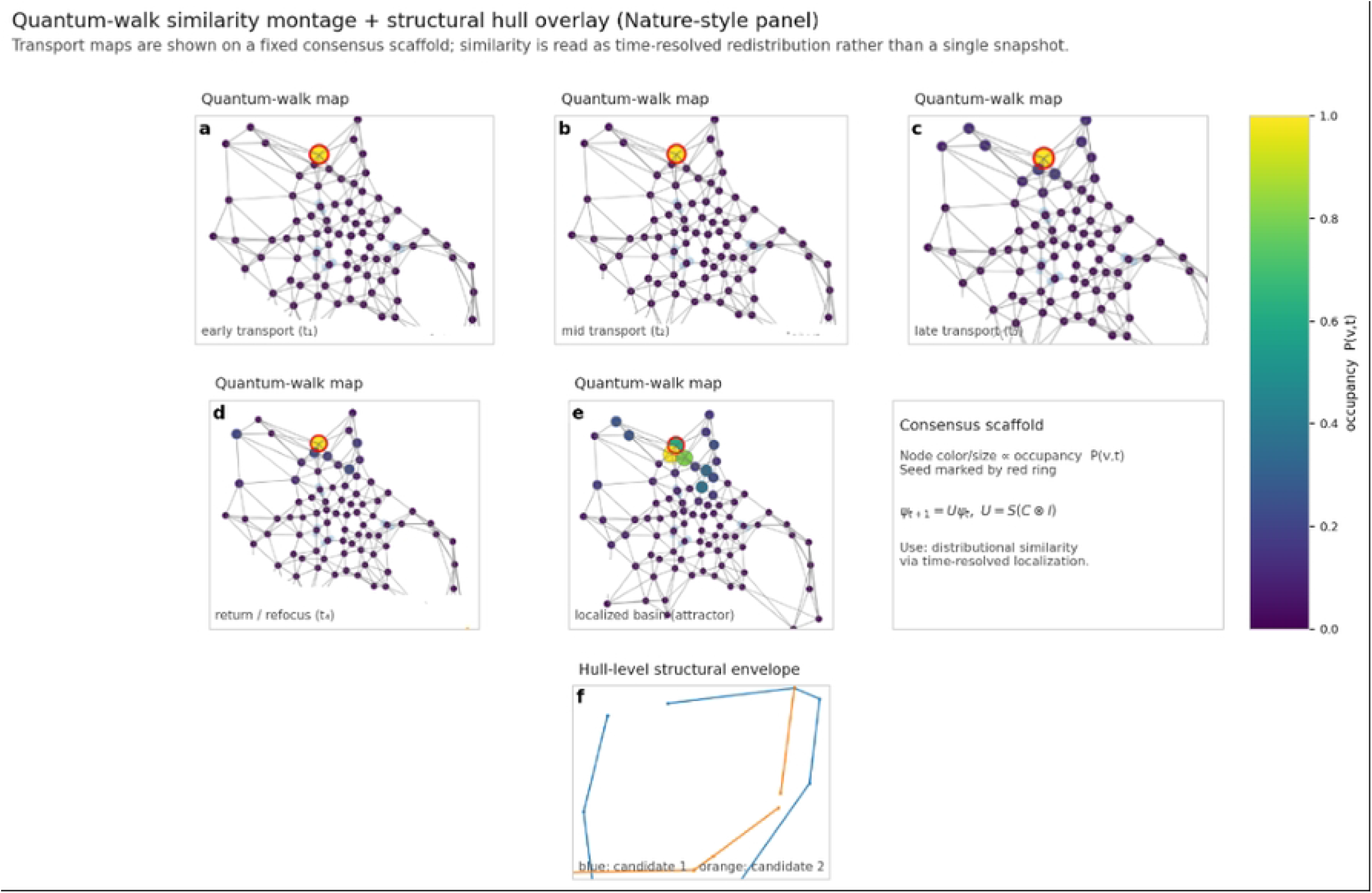
Linobectide™ “quantum superfaces” interaction-envelope map with graph-transport proxy network. The upper panel preserves the Linobectide1TM’s 2D chemical scaffold and overlays a set of semi-transparent “quantum superfaces” that act as field-like interaction envelopes. The warm red lobes concentrate around oxygen-rich regions (hydroxyl/carbonyl-associated features), blue lobes emphasize nitrogen/hetero-atom zones, and the green/purple clouds provide a broader “context” envelope across dense ring/connector regions, giving a visually intuitive map of where distinct interaction neighborhoods cluster along the molecule. Further details are provided in SI Appendix I.

**Figure X.**
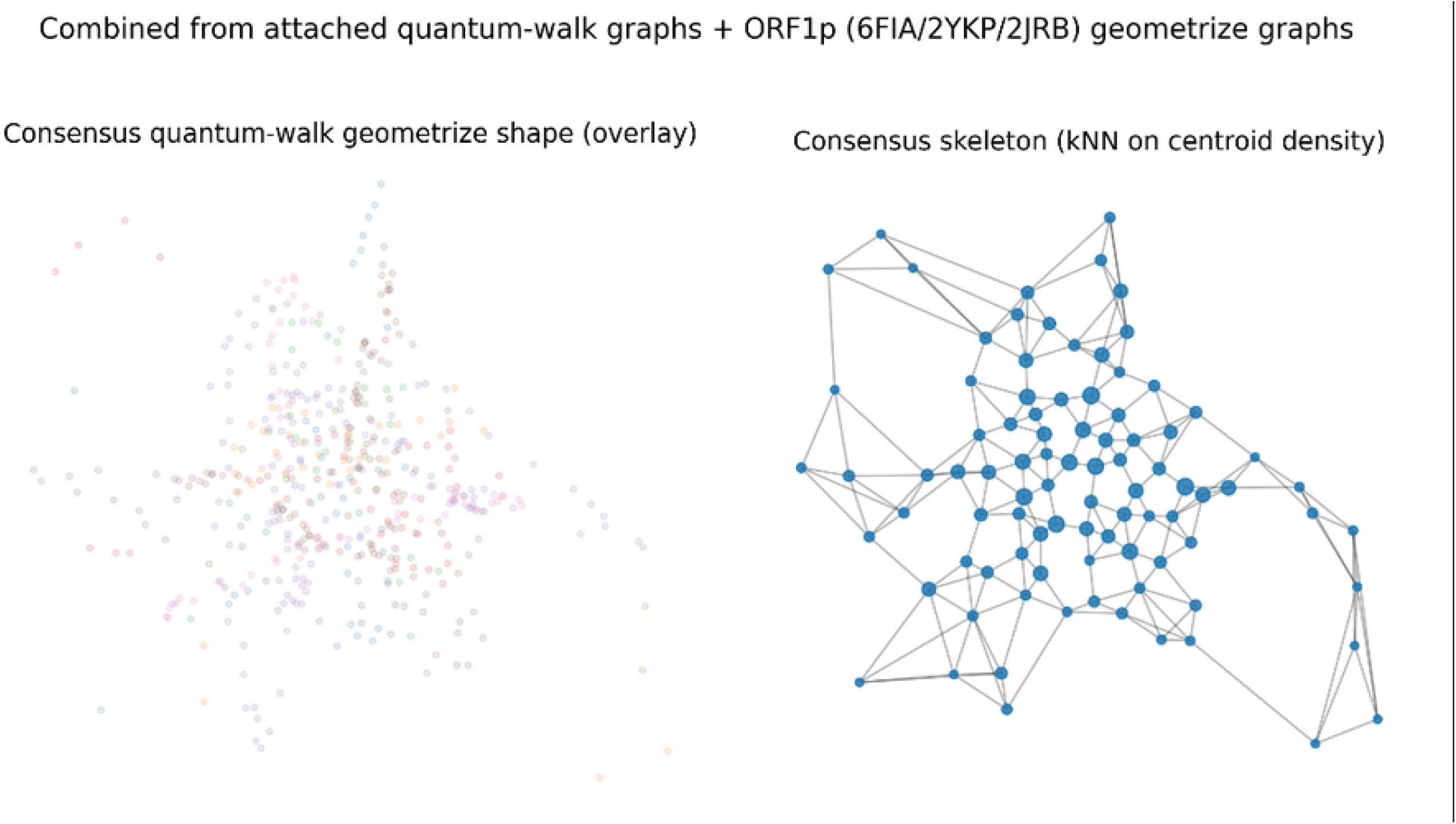
Linobectide™ composite with docking summaries, residue persistence proxies, contact-overlap structure, and graph-transport diagnostics in the 2YKP context. This composite is designed to be read as an audit-friendly “from-scores-to-structure-to-transport” story in which conventional docking outputs are first rendered as rank-ordered and distributional summaries, then unfolded into residue-resolved interaction proxies, and finally re-expressed as a quantum-walk–style transport diagnostic and a minimal circuit schematic that makes the bookkeeping assumptions explicit rather than implicit. Further details are provided in SI Appendix I.

**Figure Y.**
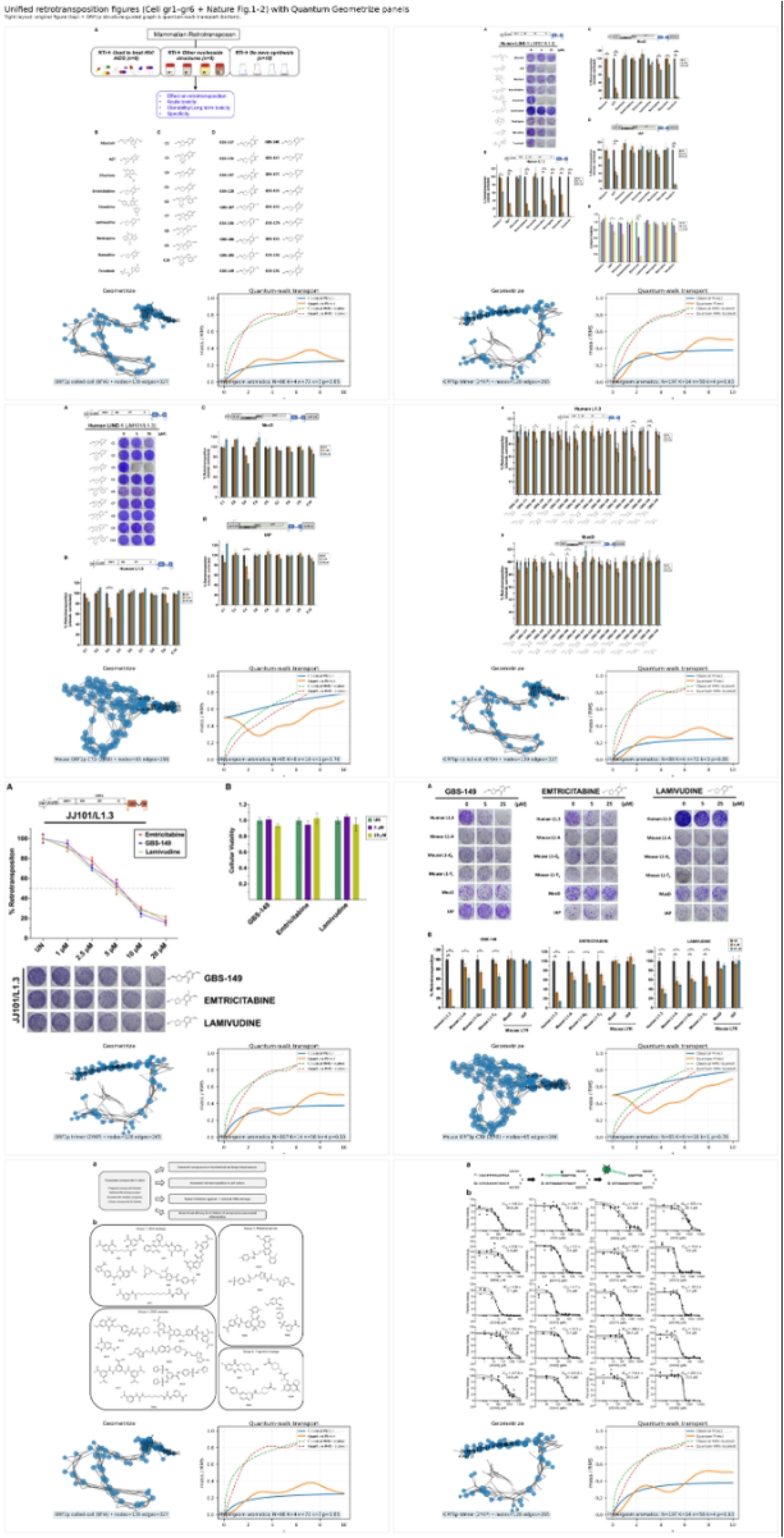
Linobectide™ unified comparative docking and persistence composite, with pocket-wise energetics, residue-frequency persistence, taxonomy-level class persistence, contact-overlap structure, and MBHA/quantum-walk scheduling schematic across 6FIA and 2YKP. This composite is intended to be read as a single, traceable “from raw docking evidence to reproducible interaction structure to method-level scheduling logic” narrative, in which the outputs of a comparative docking run are first preserved in their native provenance (a CB-Dock2 screenshot montage), then distilled into pocket-resolved energetics and residue-indexed persistence proxies that can be pooled across pockets and compared across targets, and finally connected to the workflow’s higher-level optimization control logic through explicit overlap metrics and a scheduling schematic that distinguishes diffusive and ballist Further details are provided in SI Appendix I.

**Figure Z.**
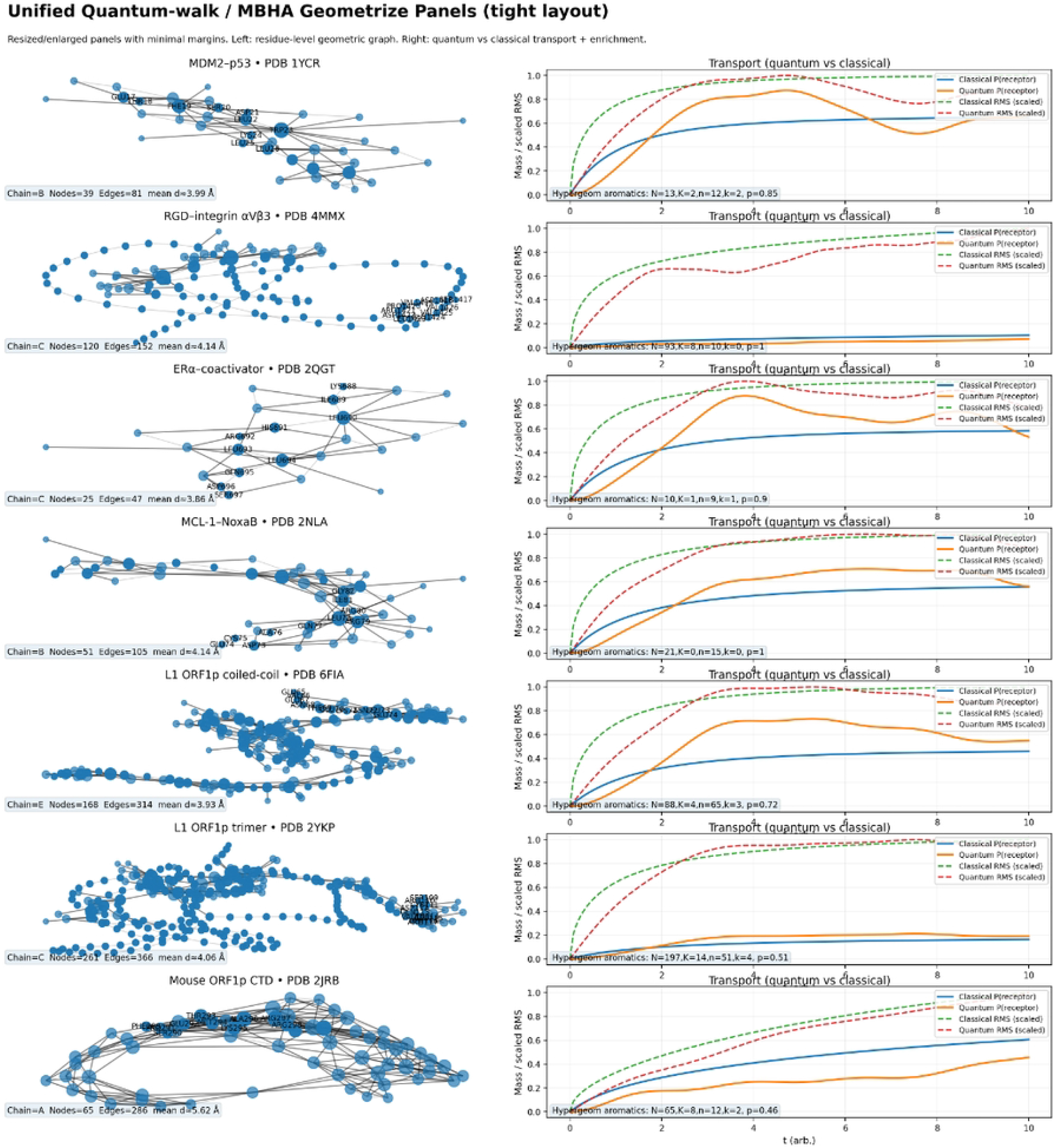
Unified docking visuals, graph-transport diagnostics, and comparative scoring for the Linobectide™ 2YKP series, with consensus-state geometry and graph-skeleton audit layer. This unified panel is constructed as a deliberately “from provenance to abstraction” figure, in which the first nine subpanels (Z.A–Z.I) preserve the primary visual artifacts of the docking and state-graph workflow, while the lower row translates those artifacts into compact, rerunnable observables— rank order, transport signature, and a cross-metric comparison—so that promotion decisions can be traced from raw evidence to interpretable summaries without collapsing the pipeline into a single scalar. Further details are provided in SI Appendix I.

**Figure W.**
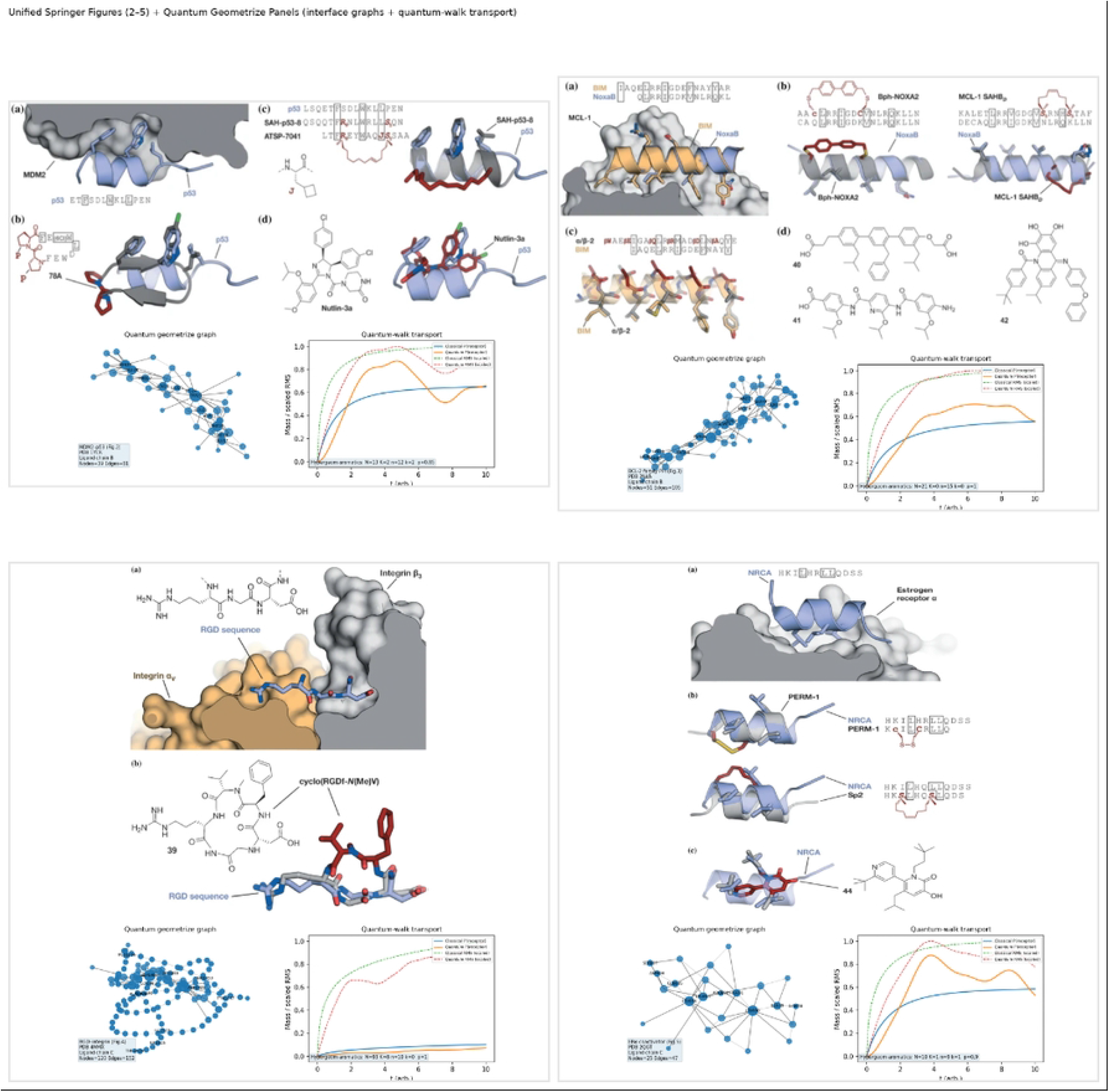
Pocket repeatability and transport-ready contact topology across state-snapshot docking annotations. Ensemble docking mosaic of Linobeqtide™–ALK complexes highlighting a conserved terminal pocket and transport-ready contact topology. This docking mosaic assembles representative Linobeqtide™–ALK complexes across distinct receptor conformations and sampling regimes, emphasizing how a shared “terminal pocket” geometry can persist even when the global fold, ligand entry path, and peripheral contacts vary. Each tile shows the receptor scaffold rendered in a dark, low-clutter style, with the ligand trace highlighted against a black field to preserve spatial readability of the binding site. Further details are provided in SI Appendix I.

**Figure S.**
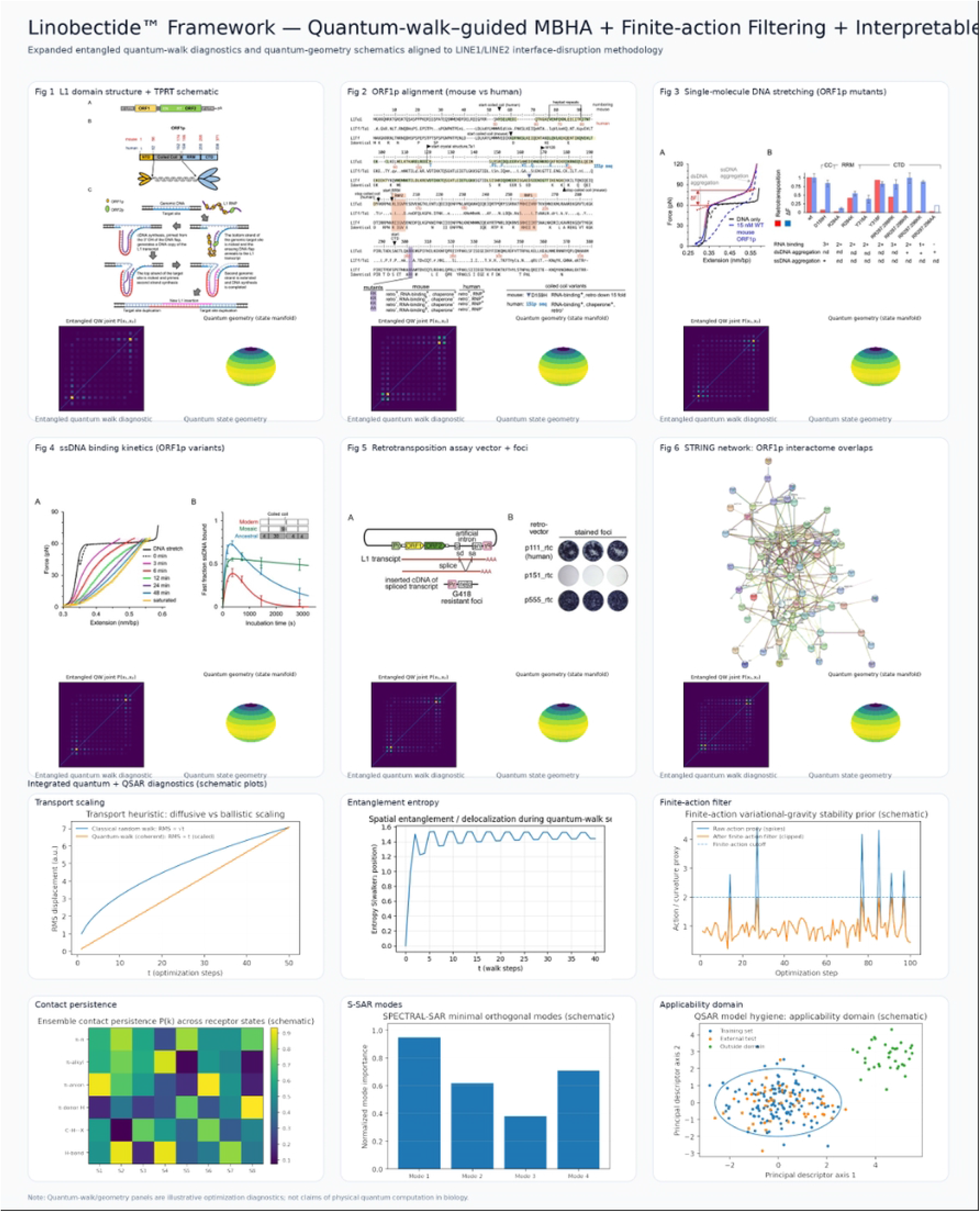
Terminal localization diagnostics and teleportation-stabilized pocket engagement for Linobectide™ candidates. This figure assembles four complementary diagnostics that translate static interaction spectra and docking energies into a transport-based picture of how each compound concentrates (or fails to concentrate) engagement onto a terminal residue subset. The analysis is explicitly framed as proxy quantum-walk machinery—an operator-driven transport model used for ranking and interpretability rather than a claim that biochemical quantum coherence is sustained in vivo. Further details are provided in SI Appendix I.

**Figure T.**
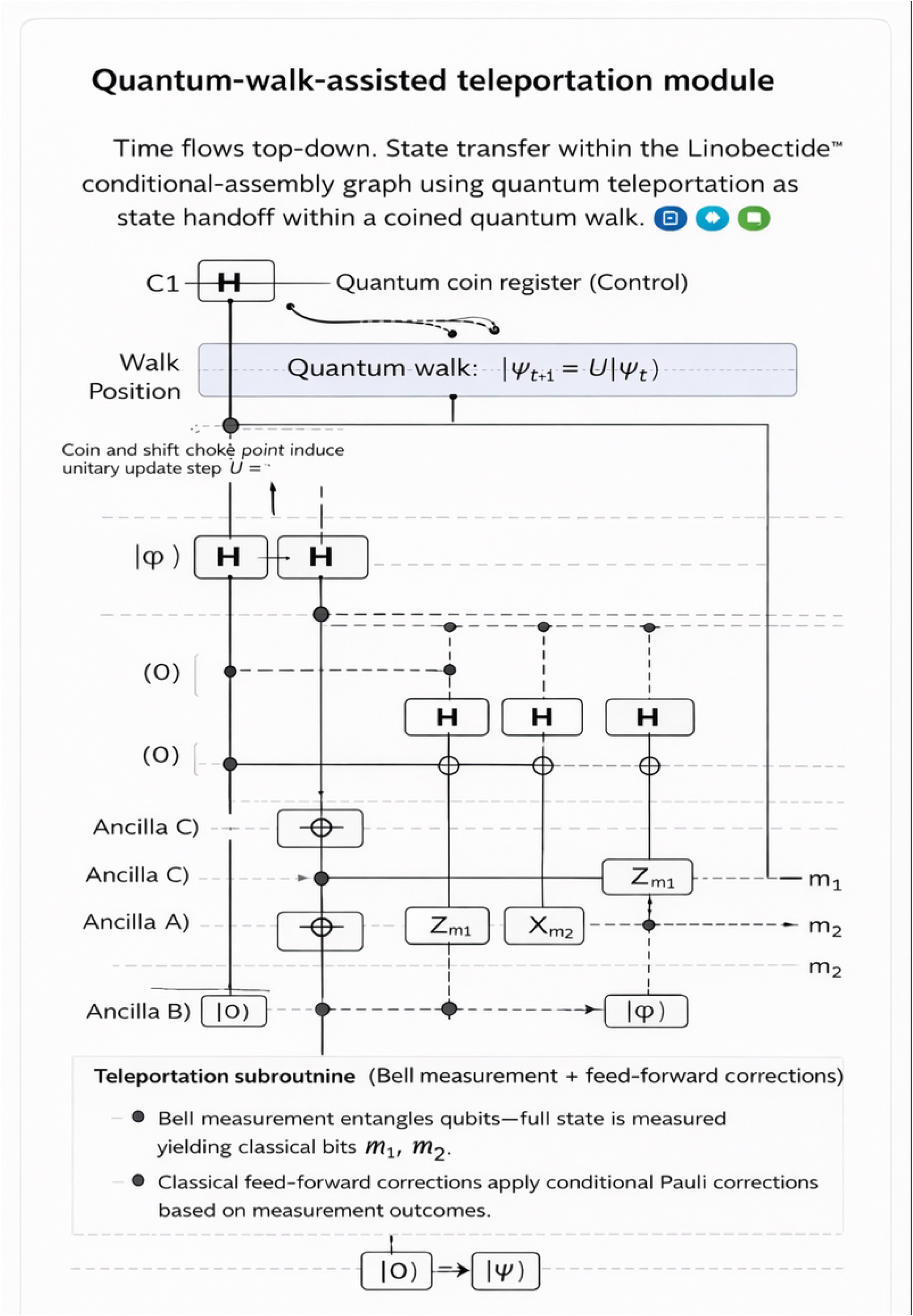
Comparative quantum docking energy landscape for Linobectide variants versus FDA/other compounds (2YKP), connecting energetic ranking to a teleportation-stabilized quantum-walk view of terminal pocket engagement. This multi-panel figure summarizes a comparative docking screen in which lower (more negative) docking energies are interpreted as stronger predicted affinity in the 2YKP pocket context. The layout is intentionally identifies the strongest binders by rank, then quantifies group-level separation, then exposes compound-level heterogeneity and tail structure, and finally consolidates the key numerical takeaways in a compact summary. Throughout, the Linobeqtide family is emphasized as a coherent reference class (blue) against a broader collection of FDA/other molecules (gray or orange). Further details are provided in SI Appendix I.

**Figure V.**
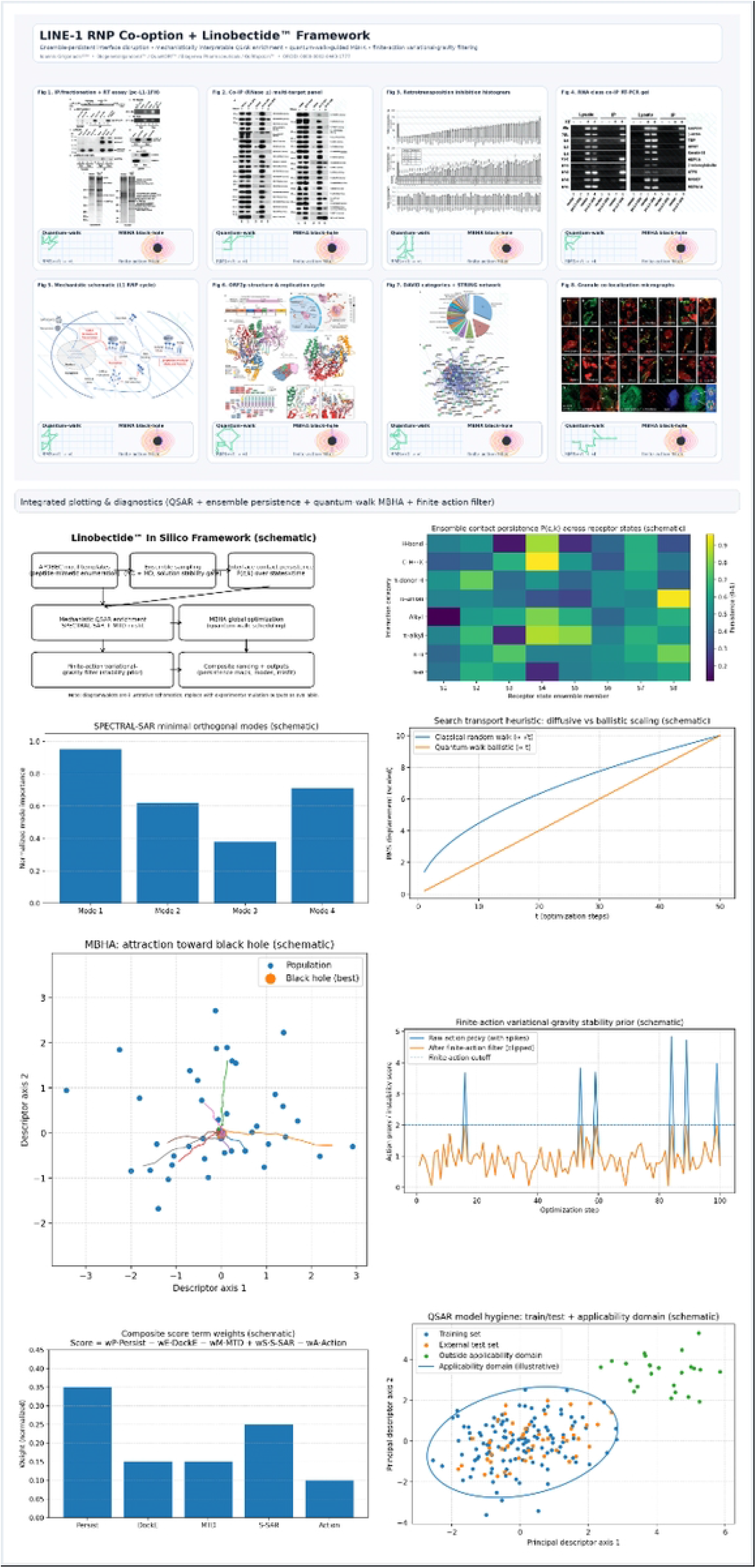
Comparative docking triage summary and strong-binding tail structure for Linobectide™ versus reference compounds. This figure condulates the “strong-binder” regime by restricting attention to negative docking energies, , thereby removing positive/near-zero outliers that otherwise dominate scale but contribute little to discriminating high-affinity candidates. Read left-to-right, the panels move from rank-ordered exemplars (who sits in the extreme tail), to distributional separation (how the cohorts differ in central tendency and spread), to a cumulative view (how often each cohort attains energies at or beyond a given threshold). Further details are provided in SI Appendix I.

## References

[1] Buzzelli M, Bianco S (2025) Uncertainty estimation in color constancy. Pattern Recognit 160:111175. 10.1016/j.patcog.2024.111175

[2] Wang ZJ, Li H, Xu ZX, Zhao SY, Wang PJ, Gao HB (2025) Adaptive learning rate algorithms based on the improved Barzilai–Borwein method. Pattern Recognit 160:111179. 10.1016/j.patcog.2024.111179

[3] Zhang Y, Hu J, Wen D, Deng W (2025) Unsupervised evaluation for out-of-distribution detection. Pattern Recognit 160:111212. 10.1016/j.patcog.2024.111212

[4] Bley F, Lapuschkin S, Samek W, Montavon G (2025) Explaining predictive uncertainty by exposing second-order effects. Pattern Recognit 160:111171. 10.1016/j.patcog.2024.111171

[5] Foucart A, Debeir O, Decaestecker C (2023) Evaluating participating methods in image analysis challenges: Lessons from MoNuSAC 2020. Pattern Recognit 141:109600. 10.1016/j.patcog.2023.109600

[6] Lo LJ, Yang CT, Chiang WC, Lin HH (2024) A quantitative method for the assessment of facial attractiveness based on transfer learning with fine-grained image classification. Pattern Recognit 145:109970. 10.1016/j.patcog.2023.109970

[7] Vovk V (2025) Conformal e-prediction. Pattern Recognit 166:111674. 10.1016/j.patcog.2025.111674

[8] Bai X, Wang X, Liu X, Liu Q, Song J, Sebe N, Kim B (2021) Explainable deep learning for efficient and robust pattern recognition: A survey of recent developments. Pattern Recognit 120:108102. 10.1016/j.patcog.2021.108102

[9] Wang J, Chen J, Zhang K, Sigal L (2024) Training feedforward neural nets in Hopfield-energy-based configuration: A two-step approach. Pattern Recognit 145:109954. 10.1016/j.patcog.2023.109954

[10] Chen K, van Laarhoven T, Marchiori E (2024) Compressing spectral kernels in Gaussian process: Enhanced generalization and interpretability. Pattern Recognit 155:110642. 10.1016/j.patcog.2024.110642

[11] Fu C, Du B, Zhang L (2024) Hybrid-context-based multi-prior entropy modeling for learned lossless image compression. Pattern Recognit 155:110632. 10.1016/j.patcog.2024.110632

[12] Hu P, Ma J, Zhang Z, Du J, Zhang J (2025) Count, decompose and correct: A new approach to handwritten Chinese character error correction. Pattern Recognit 160:111110. 10.1016/j.patcog.2024.111110

[13] Zou Y, Zhao Q, Sarker PK, Li S, Wang L, Liu W (2025) Diffusion-based framework for weakly-supervised temporal action localization. Pattern Recognit 160:111207. 10.1016/j.patcog.2024.111207

[14] Guo X, Jiang F, Chen Q, Wang Y, Sha K, Chen J (2025) Deep learning-enhanced environment perception for autonomous driving: MDNet with CSP-DarkNet53. Pattern Recognit 160:111174. 10.1016/j.patcog.2024.111174

[15] Wu Z, Sheng Z, Zhang X, Cao SY, Zhang R, Yu B, Zhang C, Yang B, Shen HL (2025) STARNet: Low-light video enhancement using spatio-temporal consistency aggregation. Pattern Recognit 160:111180. 10.1016/j.patcog.2024.111180

[16] Tao J, Chan S, Shi Z, Bai C, Chen S (2025) FocTrack: Focus attention for visual tracking. Pattern Recognit 160:111128. 10.1016/j.patcog.2024.111128

[17] Wang Y, Shao Z, Lu T, Huang X, Wang J, Zhang Z, Zuo X (2025) Lightweight remote sensing super-resolution with multi-scale graph attention network. Pattern Recognit 160:111178. 10.1016/j.patcog.2024.111178

[18] He W, Ren J, Bai R, Jiang X (2025) Two-stage rule-induction visual reasoning on RPMs with an application to video prediction. Pattern Recognit 160:111151. 10.1016/j.patcog.2024.111151

[19] dos Santos SF, Berriel R, Oliveira-Santos T, Sebe N, Almeida J (2025) Budget-aware pruning: Handling multiple domains with less parameters. Pattern Recognit 167:111714. 10.1016/j.patcog.2025.111714

[20] Khan MQ, Shahzad M, Khan SA, Fraz MM, Zhu XX (2024) Beyond local patches: Preserving global–local interactions by enhancing self-attention via 3D point cloud tokenization. Pattern Recognit 155:110712. 10.1016/j.patcog.2024.110712

[21] Wang Z, Chen L, He J, Yang L, Wang FY (2025) Exploring latent transferability of feature components. Pattern Recognit 160:111184. 10.1016/j.patcog.2024.111184

[22] Dai T, Feng Y, Chen B, Lu J, Xia ST (2022) Deep image prior based defense against adversarial examples. Pattern Recognit 122:108249. 10.1016/j.patcog.2021.108249

[23] Shabani M, Tran DT, Kanniainen J, Iosifidis A (2023) Augmented bilinear network for incremental multi-stock time-series classification. Pattern Recognit 141:109604. 10.1016/j.patcog.2023.109604

[24] Yao T, Wang R, Wang J, Li Y, Yue J, Yan L, Tian Q (2024) Efficient supervised graph embedding hashing for large-scale cross-media retrieval. Pattern Recognit 145:109934. 10.1016/j.patcog.2023.109934

[25] Liu Y, Hou X (2023) Local multi-scale feature aggregation network for real-time image dehazing. Pattern Recognit 141:109599. 10.1016/j.patcog.2023.109599

[26] Morales A, Fierrez J, Acien A, Tolosana R, Serna I (2022) SetMargin loss applied to deep keystroke biometrics with circle packing interpretation. Pattern Recognit 122:108283. 10.1016/j.patcog.2021.108283

[27] Zhang Y, Liu M, Zhang H, Sun G, He J (2023) Adaptive fusion affinity graph with noise-free online low-rank representation for natural image segmentation. Pattern Recognit 141:109611. 10.1016/j.patcog.2023.109611

[28] Fan Y, Liu J, Tang J, Liu P, Lin Y, Du Y (2024) Learning correlation information for multi-label feature selection. Pattern Recognit 145:109899. 10.1016/j.patcog.2023.109899

[29] Chen J, Song P, Zhao C (2024) Multi-scale self-supervised representation learning with temporal alignment for multi-rate time series modeling. Pattern Recognit 145:109943. 10.1016/j.patcog.2023.109943

[30] Yang Y, Hossain MZ, Stone E, Rahman S (2024) Spatial transcriptomics analysis of gene expression prediction using exemplar guided graph neural network. Pattern Recognit 145:109966. 10.1016/j.patcog.2023.109966

[31] Li C, Zhang C (2024) Toward a deeper understanding: RetNet viewed through convolution. Pattern Recognit 155:110625. 10.1016/j.patcog.2024.110625

[32] Chen L, Tang Z, Li H (2024) Improving CNN-based semantic segmentation on structurally similar data using contrastive graph convolutional networks. Pattern Recognit 155:110622. 10.1016/j.patcog.2024.110622

[33] Liu BD, Shao S, Zhao C, Xing L, Liu W, Cao W, Zhou Y (2024) Few-shot image classification via hybrid representation. Pattern Recognit 155:110640. 10.1016/j.patcog.2024.110640

[34] Akhtar M, Tanveer M, Arshad M (2024) Advancing supervised learning with the wave loss function: A robust and smooth approach. Pattern Recognit 155:110637. 10.1016/j.patcog.2024.110637

[35] Liu L, Liu Z, Chang J, Xu X (2024) A multi-modal extraction integrated model for neuropsychiatric disorders classification. Pattern Recognit 155:110646. 10.1016/j.patcog.2024.110646

[36] Song Y, Guo L, Man M, Wu Y (2024) The spiking neural network based on fMRI for speech recognition. Pattern Recognit 155:110672. 10.1016/j.patcog.2024.110672

[37] Doherty J, Gardiner B, Kerr E, Siddique N (2025) BiFPN-YOLO: One-stage object detection integrating bi-directional feature pyramid networks. Pattern Recognit 160:111209. 10.1016/j.patcog.2024.111209

[38] Cao J, Dong W, Chen J (2024) View-unaligned clustering with graph regularization. Pattern Recognit 155:110706. 10.1016/j.patcog.2024.110706

[39] Zhang J, Gao Y, Zhou M, Liu R, Cheng X, Nikolić SV, Chen S (2025) SVD-KD: SVD-based hidden layer feature extraction for knowledge distillation. Pattern Recognit 167:111721. 10.1016/j.patcog.2025.111721

[40] Pilavcı YY, Güneyi ET, Cengiz C, Vural E (2024) Graph domain adaptation with localized graph signal representations. Pattern Recognit 155:110628. 10.1016/j.patcog.2024.110628

[41] Jiang Q, Zhang Y, Bao F, Zhao X, Liu P (2022) Two-step domain adaptation for underwater image enhancement. Pattern Recognit 122:108324. 10.1016/j.patcog.2021.108324

[42] Zhong JL, Gan YF, Vong CM, Yang JX, Zhao JH, Luo JH (2022) Effective and efficient pixel-level detection for diverse video copy-move forgery types. Pattern Recognit 122:108286. 10.1016/j.patcog.2021.108286

[43] Teng Z, Cao P, Huang M, Gao Z, Wang X (2024) Multi-label borderline oversampling technique. Pattern Recognit 145:109953. 10.1016/j.patcog.2023.109953

[44] Gong J, Zhao Y, Zhao J, Zhang J, Ma G, Zheng S, Du S, Tang J (2024) Personalized recommendation via inductive spatiotemporal graph neural network. Pattern Recognit 145:109884. 10.1016/j.patcog.2023.109884

[45] Zhou F, Sun X, Dong J, Zhu XX (2023) SurroundNet: Towards effective low-light image enhancement. Pattern Recognit 141:109602. 10.1016/j.patcog.2023.109602

[46] Bayraktar E, Yigit CB (2024) Conditional-pooling for improved data transmission. Pattern Recognit 145:109978. 10.1016/j.patcog.2023.109978

[47] Hou Y, Ma Z, Liu C, Wang Z, Loy CC (2024) Network pruning via resource reallocation. Pattern Recognit 145:109886. 10.1016/j.patcog.2023.109886

[48] Zhao W, Zhao H (2024) Hierarchical long-tailed classification based on multi-granularity knowledge transfer driven by multi-scale feature fusion. Pattern Recognit 145:109842. 10.1016/j.patcog.2023.109842

[49] Liu Q, He X, Teng Q, Qing L, Chen H (2023) BDNet: A BERT-based dual-path network for text-to-image cross-modal person re-identification. Pattern Recognit 141:109636. 10.1016/j.patcog.2023.109636

[50] Cui L, Li M, Bai L, Wang Y, Li J, Wang Y, Li Z, Chen Y, Hancock ER (2024) QBER: Quantum-based entropic representations for un-attributed graphs. Pattern Recognit 145:109877. 10.1016/j.patcog.2023.109877

[51] Li Q, Yan C, Hao Q, Peng X, Liu L (2024) Graph attentive dual ensemble learning for unsupervised domain adaptation on point clouds. Pattern Recognit 155:110690. 10.1016/j.patcog.2024.110690

[52] Ji Y, Li F, Fu B, Zhou Y, Wu H, Li Y, Li X, Shi G (2024) A novel hybrid decoding neural network for EEG signal representation. Pattern Recognit 155:110726. 10.1016/j.patcog.2024.110726

[53] Yang Z, Tan Y, Yang T (2024) Large-scale multi-view clustering via matrix factorization of consensus graph. Pattern Recognit 155:110716. 10.1016/j.patcog.2024.110716

[54] Pan X, Han X, Wang C, Li Z, Song S, Huang G, Wu C (2024) A unified framework for convolution-based graph neural networks. Pattern Recognit 155:110597. 10.1016/j.patcog.2024.110597

[55] Peng L, He Y, Wang S, Song X, Dong S, Wei X, Gong Y (2024) Global self-sustaining and local inheritance for source-free unsupervised domain adaptation. Pattern Recognit 155:110679. 10.1016/j.patcog.2024.110679

[56] Yaghoobi S, Mojallali H (2016) Modified Black Hole Algorithm with Genetic Operators. International Journal of Computational Intelligence Systems 9(4):652–665. 10.1080/18756891.2016.1204114

